# Fluorescence complementation enables quantitative imaging of cell penetrating peptide-mediated protein delivery in plants including WUSCHEL transcription factor

**DOI:** 10.1101/2022.05.03.490515

**Authors:** Jeffrey W. Wang, Natalie Goh, Henry Squire, Michael Ni, Edward Lien, Eduardo González-Grandío, Markita P. Landry

## Abstract

Protein delivery to plants offers many opportunities for plant bioengineering via gene editing and through direction of protein-protein interactions. However, the delivery and confirmation of successful protein delivery to plants presents both practical and analytical challenges. We present a GFP bimolecular fluorescence complementation-based tool, delivered complementation *in planta* (DCIP), which allows for unambiguous and quantitative measurement of protein delivery in leaves. Using DCIP, we demonstrate cell-penetrating peptide mediated cytosolic delivery of peptides and recombinant proteins in *Nicotiana benthamiana*. We show that DCIP enables measurement of delivery efficiency and enables functional screening of cell penetrating peptide efficacies for in-plant protein delivery. Finally, we demonstrate that DCIP detects cell penetrating peptide mediated delivery of recombinantly expressed proteins such as mCherry and Lifeact into intact leaves. Finally, we also demonstrate, for the first time, delivery of a recombinant plant transcription factor, WUSCHEL (AtWUS), in *N. benthamiana*. RT-qPCR analysis of AtWUS delivery in *Arabidopsis* seedlings also suggests delivered WUS can recapitulate AtWUS-overexpression transcriptional changes. All combined, DCIP offers a new and powerful tool for interrogating cytosolic delivery of proteins in plants and highlights future avenues for engineering plant physiology.

## Introduction

The delivery of proteins to walled plant cells remains an ongoing challenge. In addition to the cell membrane, the plant cell wall is an effective cellular barrier not only to naturally occurring pathogens but also to introduced macro-biomolecules: DNA, RNA, and proteins. While several tools exist for the delivery of nucleic acids in plants, very few enable delivery of proteins to walled plant cells. The development of CRISPR-Cas9 (Jinek, Chylinski et al. 2012) and other DNA editing tools (Li, Li et al. 2020) has only increased the need for working protein delivery tools, which could accelerate basic research, spawn novel agricultural biologic agents, or potentiate DNA-free gene editing of plants. Recent discoveries in small microproteins (Dolde, Rodrigues et al. 2018) that modulate plant growth evinces a new class of possible protein cargoes if these proteins could be delivered into leaves. These motivations have led researchers to develop novel nanoparticle-based strategies for the delivery of biomacromolecules to walled plant cells. For example, multiple technologies have been developed to deliver siRNA to plants using diverse vehicles such as single walled carbon nanotubes (Demirer, Zhang et al. 2020), DNA nanostructures (Zhang, Demirer et al. 2019), carbon dots (Schwartz, Hendrix et al. 2020), and gold nanoparticles (Zhang, Goh et al. 2022). Fewer have demonstrated delivery of proteins using cell penetrating peptides (Chang, Chou et al. 2007, Guo, Itami et al. 2019). Despite these proof-of-principle advances, protein delivery to walled plant cells remains largely dependent on biolistic delivery, which requires protein dehydration (and thus potential inactivation) to a gold particle surface and forceful and injurious rupture of plant membranes to accomplish delivery in a low throughput and low efficiency manner (Hamada, Liu et al. 2018). One main barrier to the use of nanotechnologies for plant biomolecule delivery, and specifically cell penetrating peptides for protein delivery to plants, is the lack of quantitative validation of successful intracellular protein delivery. This barrier makes it difficult to unilaterally distinguish successful protein delivery from artefact, lytic sequestration, or quantitatively compare delivery efficiency of different peptides (Wang, Cunningham et al. 2021).

This lack of tools to quantify successful protein delivery in plants is due to the near universal dependence of confocal microscopy to validate delivery of fluorescent proxy cargoes. However, confocal microscopy in plant tissues pose a set of unique problems that make it challenging to distinguish artefact from signal and make absolute quantification of signal impossible. Aerial tissues of plants are heterogeneous, highly light scattering, and possess intrinsic auto-fluorescence (Donaldson 2020), which makes it difficult to distinguish signal from noise. Furthermore, unlike mammalian cells, the plant cell cytosol is highly compressed against the cell wall by the plant’s large central vacuole, making imaging of cytosolic contents challenging due to the small surface area of cytosolic contents (Serna 2005). In addition, the plant cell is surrounded by a porous and adsorbent cellulosic wall that is 100-500nm thick (Sugiura, Terashima et al. 2020) which spans the Rayleigh diffraction resolution limit of visible light imaging and the axial resolution of most confocal microscopes. Together, the small cytosolic volume which is proximal to the cell wall makes it impossible to distinguish – with the necessary spatial precision – the location of fluorescent cargoes near versus imbedded in the cell wall, or inside the cell cytosol (Zhang, Goh et al. 2022), without super-resolution microscopy (Pawley 2006). Additionally, free fluorophore from cargo degradation (Lacroix, Vengut-Climent et al. 2019) or endosomal entrapment of cargoes would contribute to measured fluorescence intensity and intracellular colocalization in plants but fail to correlate with successful delivery. For these reasons, gauging cellular uptake of cargoes based solely on confocal microscopy data of fluorophore-tagged cargo in plants does not confirm successful intracellular delivery nor provide quantitative data for effective uptake. These barriers have made biomacromolecule delivery in plants, particularly protein delivery, exceptionally challenging. We therefore developed a versatile, unambiguous platform to confirm the delivery of proteins in walled plant tissues and demonstrate it is possible to quantify protein delivery efficiency across different protein sizes.

We designed an *Agrobacterium tumefaciens* expression mediated, GFP-complementation based, red/green ratiometric sensor for the detection of protein delivery in plants using confocal microscopy (DCIP, **D**elivered **C**omplementation ***i****n **p**lanta*). In this technique, sfGFP is split between a larger non-fluorescent fragment (sfGFP1-10) and a smaller peptide strand (GFP11) (Hu, Chinenov et al. 2002, Kamiyama, Sekine et al. 2016). When GFP11 is delivered to the same compartment as sfGFP1-10, only then is GFP fluorescence is reconstituted (Figure 1, A). This method has the critical benefit of only producing signal if the peptide tag remains intact, is successfully delivered to the cytosol, and is not sequestered in lytic organelles or trapped in the apoplast. GFP11 also serves as an excellent reporter tag because its short length (16AA) is accessible to chemical synthesis and because it is readily incorporated into recombinant proteins as a terminal tag. The final design of DCIP was also guided by the desire to perform automated image analysis within complex leaf tissues and thus reduce the risk of bias during analysis.

**Figure 1.**
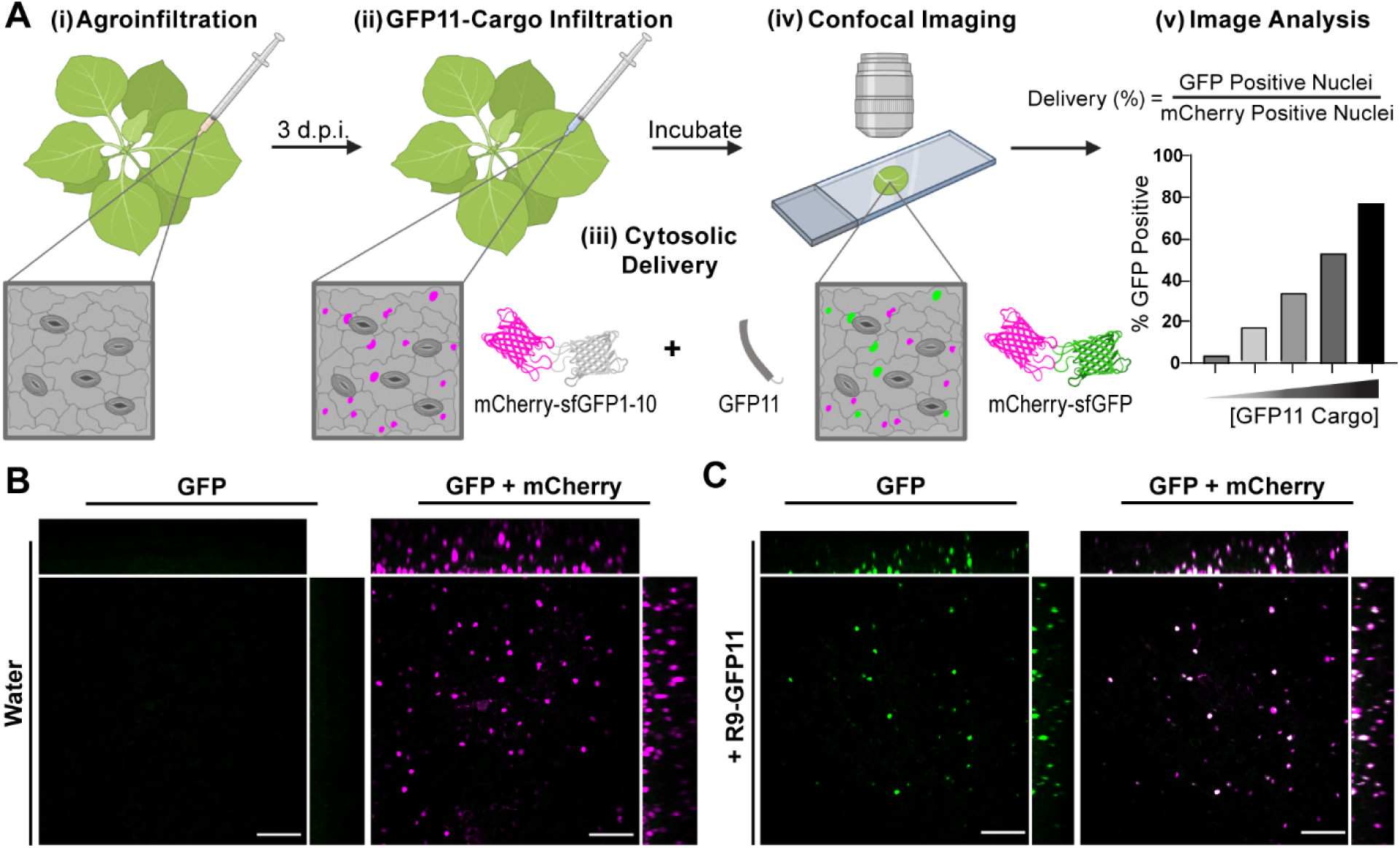
Demonstration of DCIP for peptide delivery. A, Workflow schematic for DCIP. (i) *N. benthamiana* are agroinfiltrated with the DCIP vector. (ii) 3 d.p.i. plants are infiltrated with the peptide or protein cargo fused to GFP11. (iii) During incubation, GFP11 is internalized into plant cells and if cytosolic delivery occurs, GFP11 is able to complement GFP1-10 and sfGFP fluorescence is recovered. (iv) Post-incubation, leaf discs are imaged and analyzed using Cell Profiler by using mCherry fluorescence to identify cells via their fluorescent nuclei. (v) The sfGFP fluorescence is normalized to mCherry fluorescence to account for variability in DCIP expression and the number of GFP positive cells relative to the total number of mCherry positive cells is determined as an analog to delivery efficiency. B, Representative maximum intensity projection of a leaf disc expressing DCIP infiltrated with water. sfGFP fluorescence is pseudocolored green (left) and two-color overlay with mCherry fluorescence, pseudocolored magenta, resulting in a white appearance after overlay (right). mCherry expressing cells possess nuclei presenting as small, round fluorescent bodies amenable to automated image analysis. Orthogonal projections demonstrate depth of imaging in leaves. C, Equivalent images of DCIP expressing leaf after treatment with 100 µM R9-GFP11 for 4 hours and showing that delivery capability extends throughout the full thickness of the leaf tissue. Scale bar represents 100 µm.

While bimolecular fluorescence complementation and nanoluciferase complementation have been previously applied for measuring cell penetrating peptide (CPP) mediated delivery in mammalian cells (Milech, Longville et al. 2015, Schmidt, Adjobo-Hermans et al. 2015, Teo, Rennick et al. 2021), it has not been employed to confirm the delivery of bio-cargoes to plant cells. CPPs, are small cationic or amphipathic peptides that when conjugated to cargoes, enable cytosolic delivery (Langel 2011). We utilized CPPs to test DCIP due to their synthetic accessibility, previous deployment in plant-tailored delivery schemes (Numata, Horii et al. 2018, Thagun, Horii et al. 2022), and because much of their underlying cell penetrating mechanisms in plants remain unstudied. In doing so, we show that DCIP can confirm the delivery of as little as 10µM of GFP11 peptide via confocal imaging and that nearly comprehensive (>78%) delivery to leaf cells is achieved with 300 µM of CPP delivered GFP11. Furthermore, we used DCIP to quantify the relative effectiveness of several popular CPPs (TAT, R9, BP100) to accomplish protein delivery in plants, and reveal that R9 mediated delivery in leaves is largely independent of endocytosis. We also used DCIP to probe the stability of R9-GFP11 in leaf tissues, and show a concomitant disappearance of tissue-localized GFP11 and GFP complementation signal within 24 hours. Finally, we utilize DCIP to demonstrate CPP-mediated delivery of several recombinant proteins into the cytosol of mesophyll and pavement cells. As a proof of concept, we also use DCIP to show that CPP based delivery can be used to form novel protein-protein interactions in live plants and demonstrate the delivery of *Arabidopsis* WUSCHEL transcription factor, evincing the possibility of physiologic engineering of plants using delivered proteins or peptides.

## Results

### Design of the delivered complementation *in planta* (DCIP) sensor system and workflow

Although GFP bimolecular fluorescence complementation has been used to detect CPP-mediated delivery in mammalian cells, it has not yet been employed in plants; where it is arguably most beneficial to confirm successful biomolecule delivery. We therefore developed imaging-based strategy for quantifying delivery mediated GFP complementation with DCIP. Advantages of an image-based approach include the ability to assess delivery in the complex structure of leaves and the removal of ambiguity caused by total lysis as required by a luciferase-complementation based approach (Teo, Rennick et al. 2021).

The delivery sensor protein (DCIP) consists of sfGFP1-10 C-terminally fused to mCherry with an N-terminal SV40 NLS (Hicks, Harley et al. 1995) and is transiently expressed in leaves using agrobacterium (Supplemental Figure S1). For DCIP, we chose a mCherry fusion for three reasons: (1) mCherry is easy to spectrally resolve from plant autofluorescence (2) a constitutive fusion allows identification of positively *A. tumefasciens* transfected cells and (3) mCherry fusion permits ratiometric quantification of GFP bimolecular fluorescence complementation since the relative expression of sfGFP1-10 is tied to the expression of mCherry by direct fusion. Because plant cells are heterogeneous in shape and have many autofluorescent bodies, we localized the sensor to the nucleus to produce a round, uniform object that is amenable to automated image analysis and provides unambiguous confirmation of successful delivery of GFP11 or GFP11-tagged cargoes. We also hypothesized that the NLS localization of DCIP should allow the detection of a broad range of sizes of delivered cargoes as the size exclusion limit for efficient transport through the nuclear envelop is presumed to be greater than 60 kDa (Wang and Brattain 2007). We constructed the coding sequence of DCIP sequence by traditional restriction ligation cloning and the final transcriptional unit assembly was performed using Goldenbraid 2.0 (Sarrion-Perdigones, Vazquez-Vilar et al. 2013). In tandem, we also developed a cytosolically localized version of DCIP, cytoDCIP, which lacks SV40 NLS. These constructs were transformed into *A. tumefaciens* and agroinfiltrated in *Nicotiana benthamiana* plants. *N. benthamiana* was chosen as a model plant due to its common use in transient expression experiments as well as in delivery experiments (Martin, Kopperud et al. 2009). Successful expression of intact DCIP three days post agroinfiltration (d.p.i.) was verified by microscopic observation (Figure 1, B) and by Western blot using an anti-mCherry antibody (Supplemental Figure S2), and prior to attempts at GFP11 or GFP11-tagged cargo delivery.

A typical experimental workflow using DCIP is provided in Figure 1, A. The DCIP protocol involves transient expression of DCIP in *N. benthamiana*. 3 d.p.i., leaves are infiltrated with an aqueous solution of cargo that contains the GFP11 tag. Immediately after infiltration, the infiltrated leaves are either left intact or a leaf disc is excised from the infiltrated area and plated on pH 5.7 ½ MS. After a predetermined incubation time, the leaves are imaged on a confocal laser scanning microscope. For quantitative imaging, the resulting images are then automatically analyzed using Cell Profiler (Stirling, Swain-Bowden et al. 2021) for nuclear sfGFP and mCherry fluorescence (Figure 1, A). A detailed methodology is provided in the Methods section.

For a first proof of concept, we used the nona-arginine (R9) cell penetrating peptide fused to GFP11 by a (GS)_2_ linker to validate the functionality of DCIP to detect successful GFP11 delivery. R9 was chosen due to its known effectiveness in both plant (Numata, Horii et al. 2018) and mammalian (Kosuge, Takeuchi et al. 2008) systems as well as its relatively well characterized mechanism of action in mammalian cells (Wallbrecher, Ackels et al. 2017). Without infiltration or infiltration with water, no sfGFP fluorescence is observed (Figure 1, A left) and only mCherry containing nuclei can be seen (right). Upon infiltration with 100 μM R9-GFP11, we observed robust sfGFP complementation at 4-5H post-infiltration that colocalized with mCherry (Figure 1, B). The timing of 4-5H was determined by balancing the reported mammalian uptake kinetics for R9 peptides (<1H) (Kosuge, Takeuchi et al. 2008, Brock 2014) against the relatively slow process of GFP complementation which required >5H for total complementation in our hands using an *in vitro* system with recombinant sfGFP1-10 (Supplemental Figure S3, A). For comparison, previous dye labeled cargo CPP-mediated delivery experiments in plants often used time points of about 2H (Numata, Horii et al. 2018). In this case, an 8mm leaf disc was excised from peptide infiltrated tissue and plated onto ½ MS to control possible apoplastic flow and uncontrolled drying of the infiltrated liquid which may change the effective concentration of R9-GFP11 the cells experience. An orthogonal projection (Figure 1, B) shows GFP complementation deep (∼100 μm) into the z-axis of the leaf in both pavement cells and mesophyll cells. Imaging at a lower magnification shows efficient delivery throughout the leaf disc using DCIP (Supplemental Figure S4).

### Validating DCIP to quantify protein delivery efficiency using R9-GFP11

We next assessed whether DCIP would be able to quantify the relative effectiveness of peptide delivery *in planta*. We once again used R9-GFP11 for validation experiments and infiltrated a range of concentrations from 0-100 μM R9-GFP11 into DCIP expressing *N. benthamiana*. The initial concentration range was determined from previously reported effective concentrations for mammalian cells (Pantarotto, Briand et al. 2004, Kosuge, Takeuchi et al. 2008). After 4-5 hours of incubation, we observe a clear concentration dependent upshift of the green/red ratio in DCIP expressing nuclei (Figure 2, A). Five experimental repeats (one plant per repeat) were acquired using this methodology. The resulting mean green/red ratio from each repeat were averaged and show a statistically significant increase of green/red at concentrations greater than or equal to 50 μM R9-GFP11 (Figure 2, B). Further processing of data using a percent GFP positive nuclei approach to account for variations in quantitating fluorescence data in microscopy also show similar trends (Figure 2, C left). We also observe slightly improved linearity of response with respect to R9-GFP11 concentration using percent positive instead of averaged green/red ratio. We then defined delivery efficiency by normalization of the percentage of GFP positive cells with respect to the 100 μM treatment (Figure 2, C right). Normalization helps reduce the biological variation in the absolute uptake efficiency we observed (Supplemental Figure S5). We attribute this lack of lower-end sensitivity to erroneous detection of autofluorescent bodies in the 0 μM control used for thresholding. From this initial performance study, we observed that relatively higher concentrations of R9-cargo are required for efficient delivery when compared to mammalian cells which can undergo delivery at low micromolar concentrations (Schmidt, Adjobo-Hermans et al. 2015). However, we also observed that relatively high delivery efficiency (>40% positive) could be achieved in plants when leaves were treated with 100 μM R9-GFP11 and greater (Figure 2, C).

**Figure 2.**
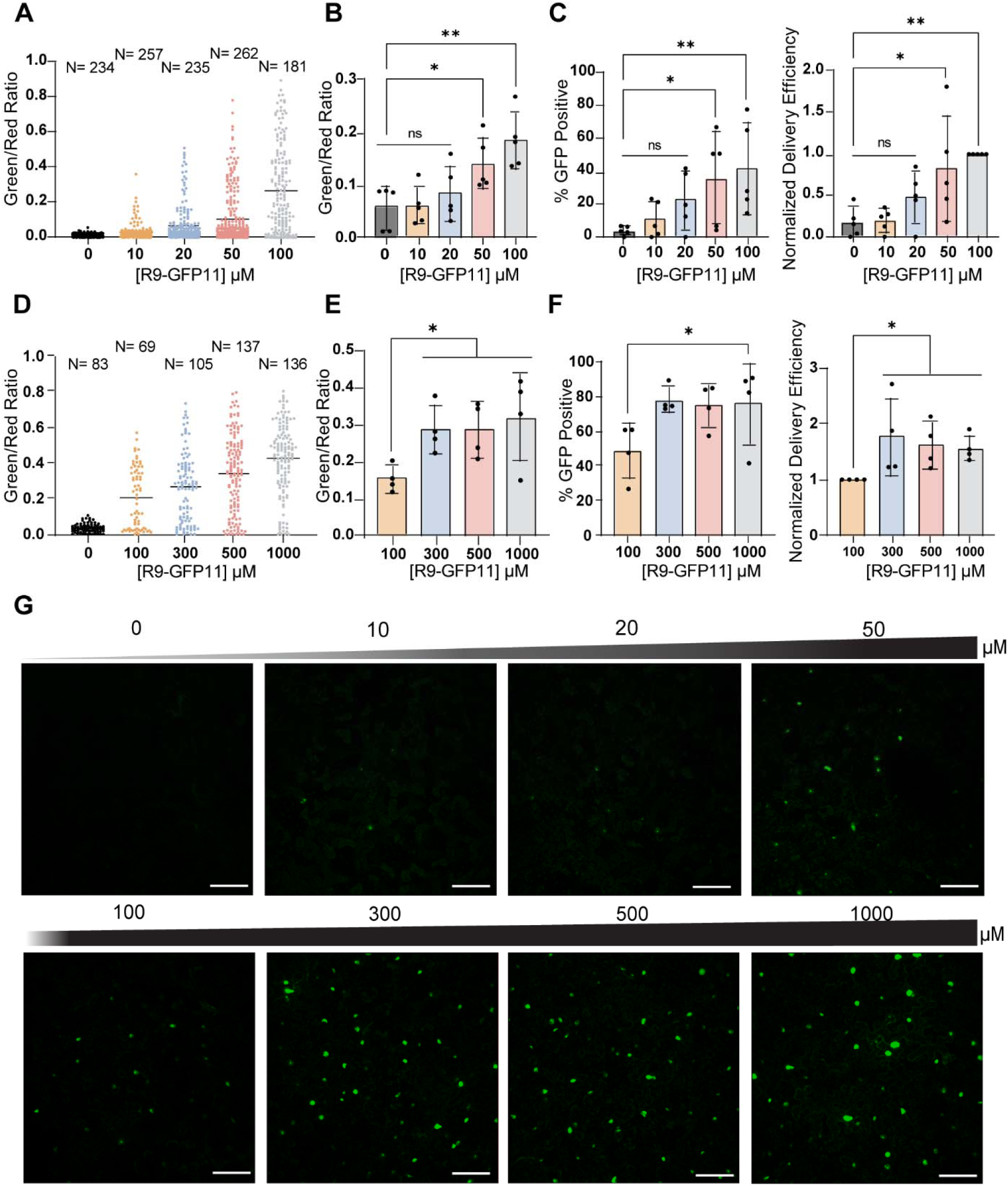
Validation of DCIP using concentration titrations of R9-GFP11. A, Representative green/red ratio in *N. benthamiana* expressing DCIP infiltrated with 0-100 µM R9-GFP11 or a water control for 4-5H. Each point represents the relative fluorescence from sfGFP caused by delivered complementation and mCherry expression in a single nucleus. Successful delivery of the GFP11 cargo results in an increase of the green/red ratio. B, Mean green/red ratio averaged across five plants as experimental repeats (N=5). Error bars represent standard deviation of the group repeats. Statistical comparisons between each treatment condition and the non-treated control with a Kruskal-Wallis test in combination with Dunn’s multiple comparison’s test. C, Percentage of GFP positive cells averaged across N=5 experimental repeats and corresponding delivery efficiency normalized to the 100μM treatment group. D, Representative green/red ratio after treating leaf discs with 100-1000 µM R9-GFP11 for 4-5H. E, Mean green/red ratio averaged across four experimental repeats (N=4) after treating leaf discs with 100-1000 µM R9-GFP11 for 4-5H. F, Average percent GFP positive cells and calculated normalized delivery efficiency of leaves treated with 100-1000 µM R9-GFP11 for 4-5 hours (N=4). * denotes p<0.05, ** p<0.01, and ns p>0.05. G, Representative single color maximum intensity projections of leaves infiltrated with 0-1000 µM R9-GFP11 and incubated for 4-5 hours. Nuclei exhibiting delivered complementation appear as round, green objects. Scale bar is 100 µm.

After a low concentration-range validation, we sought to determine at what point the DCIP signal saturates, representing the maximal possible protein delivery efficiency in plant leaves. With the aforementioned workflow, DCIP expressing leaves were infiltrated with 0-1000 μM R9-GFP11 in water and incubated as leaf discs for 4-5H. Once again, we observe strong upshifting of the green/red ratio of DCIP nuclei in treated leaves as a function of R9-GFP11 concentration (Figure 2, D). After four experimental repeats (one plant per repeat), we also observe that concentrations greater than 300 μM R9-GFP11 show a statistically significant increase in delivery compared to 100 μM R9-GFP11 and a relative saturation in green/red ratio at concentrations at 300 μM and higher (Figure 2, E). Similarly, the data analyzed using percent GFP positive cells show a saturation at about 75% positive at 300 μM and above, while the difference is only significant if the delivery efficiency is normalized to the 100 μM control (Figure 2, F). Examples of maximum intensity projections of the GFP channel for a titration DCIP experiment are provided in Fig 2g. Importantly, although quantitation failed to detect statistical significance across samples treated with less than 50 μM R9-GFP11, successful delivery could still be occasionally observed for these low peptide concentrations as sfGFP complemented nuclei (Figure 2, G). Multi-channel images showing mCherry expression show strong colocalization of the sfGFP signal and mCherry nuclear signals at all tested concentrations (Supplemental Figure S6).

### Assessing CPP performance and mechanism with DCIP

After validating DCIP for quantitative peptide delivery, we assessed whether DCIP could be used to screen for effective cell penetrating peptide sequences in leaves. Three commonly used cell penetrating peptide sequences were assessed (Figure 3, A). BP100 is a microbially derived CPP that has been previously reported to be effective in plants through a dye conjugation and delivery experiment (Numata, Horii et al. 2018). TAT is an arginine-rich HIV-1 derived peptide and one of the first cell penetrating peptides characterized (Ziegler, Nervi et al. 2005). R9 is a derivative of TAT where all amino acids are substituted for arginine (Kosuge, Takeuchi et al. 2008). Each of these CPPs were produced through solid-phase synthesis as fusions to GFP11, separated by a short (GS)_2_ linker. We used an *in vitro* bimolecular fluorescence complementation assay to ensure that the CPP fusions did not interfere with complementation activity (Supplemental Figure S3). After infiltrating 100 μM of each peptide construct into DCIP expressing leaves and incubating for 4-5H, confocal image analysis revealed that both TAT and R9 were effective at delivering GFP11 into plant cells and enabled delivery efficiencies ranging from 30-80% (Figure 3, B). R9 appeared to be the most effective of the tested CPPs with TAT being 0.77 times as effective as R9, and BP100 or GFP11 alone showing no statistically significant signal. These results support that without a CPP, GFP11 is not able to enter the cytosol of plant cells. To our surprise, BP100 mediated delivery was not statistically significantly better than either the water infiltration control or GFP11 alone. Once again, although BP100 was not statistically significantly better than the water control or GFP11 alone, we were able to observe rare instances of successful delivery for 100 μM BP100-GFP11 (Supplemental Figure S7) but not for the negative control. Closer inspection of the imaged nuclei also revealed strong nucleolar localization of sfGFP in TAT-GFP11 and R9-GFP11 treatments (Figure 3, C). These images suggest that R9 and TAT remain intact when bound to sfGFP1-10 in the cell, as poly-arginine motifs are known to localize to the nucleolus (Martin, Ter-Avetisyan et al. 2015).

**Figure 3.**
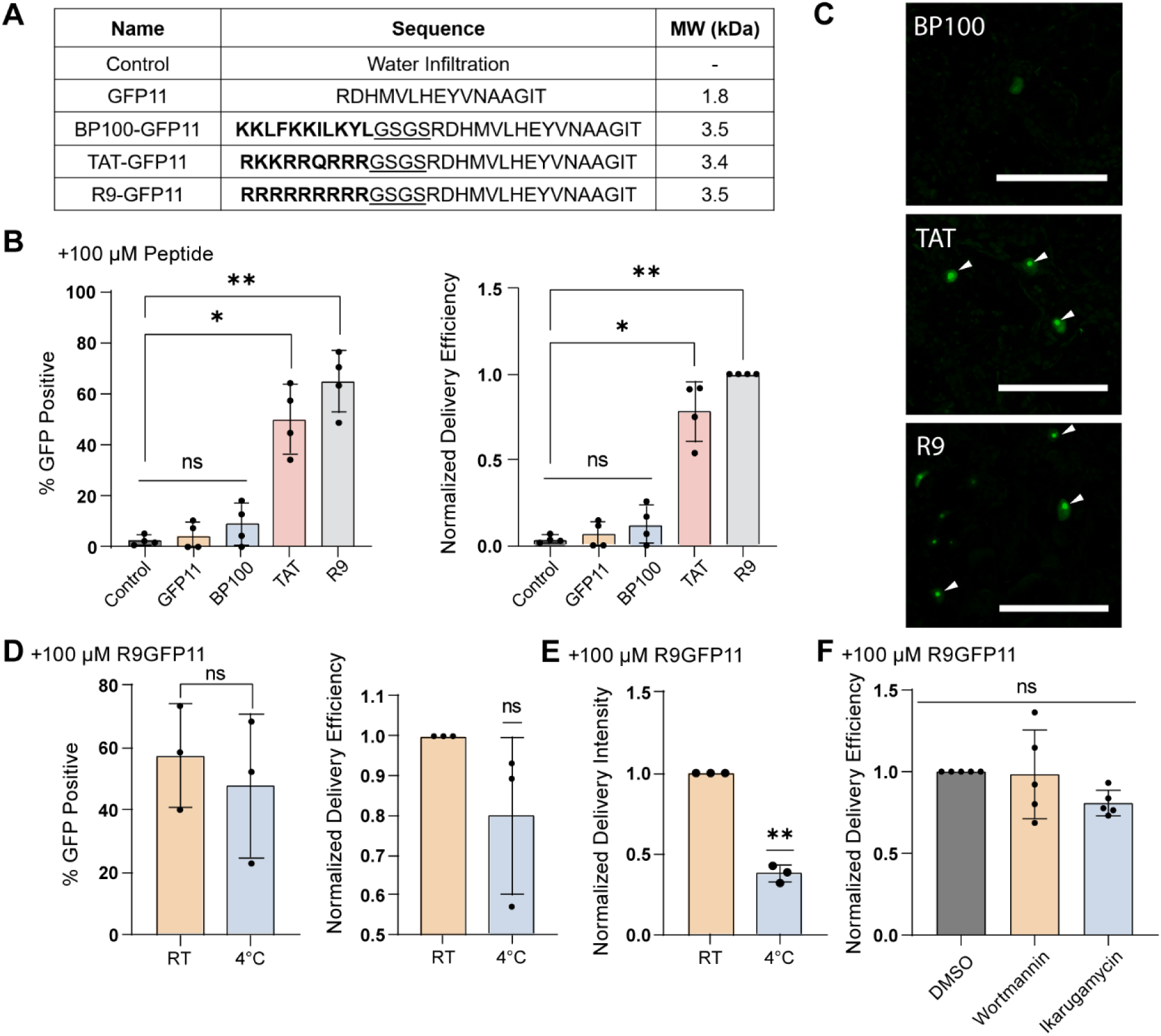
Investigating CPP performance using DCIP. A, Peptides tested for delivery of GFP11 and their corresponding sequence and molecular weights (kDa) with a water infiltration negative control. CPP sequences are bolded and the flexible GS linker is underlined. B, Percent GFP positive nuclei and normalized delivery efficiency of leaf discs incubated with 100µM with varying CPP-GFP11 conjugates for 4-5H (N=4). Normalization with respect to the 100µM R9-GFP11 group. C, Representative micrographs of sfGFP fluorescent nuclei as the result of successful GFP11 delivery. White arrows point toward enhanced nucleolar localization of DCIP after delivery using arginine-rich CPP. D, Average percent GFP positive and normalized delivery efficiency in leaf discs treated with 100µM R9-GFP11 and either left at room temperature or kept at 4°C for 4-5H (N=3). E, Average normalized delivery intensity in leaf discs treated with 100µM R9-GFP11 and either left at room temperature or kept at 4°C for 4-5H (N=3). Delivery intensity is calculated by normalizing the mean green/red ratio of the 4°C treatment to that of the room temperature treatment. For normalized results of the low-temperature treatment, a one-sample t-test comparing to the ideal value of 1.0 was used. F, Normalized delivery efficiency in leaves infiltrated with 100µM R9-GFP11 and co-infiltrated with either DMSO, 10µM ikarugamycin, or 40 µM wortmannin for 4-5H. Unless otherwise indicated, Kruskal-Wallis test followed by Dunn’s multiple comparisons test was performed for all statistical comparisons where ns = p>0.05, * = 0.01<p<0.05, and ** = p<0.01. Scale bar is 100 µm.

After validation of R9-GFP11 as the best performing CPP for protein delivery in plants, we sought to probe the mechanism by which R9 delivers cargoes to the plant cell. Specifically, we probed whether R9 delivery was endocytosis dependent or cell autonomous. The delivery efficiency (based on percentage of positive cells) in leaf discs infiltrated with 100 μM R9GFP11 and incubated at 4°C or room temperature was not statistically different. However, the normalized delivery intensity, as defined by green/red ratio normalized to the room temperature treatment showed that the 4°C discs possessed lower sfGFP fluorescence. This aligns with our *in vitro* data showing that bimolecular fluorescence complementation is possible at 4°C although somewhat compromised in efficiency (Supplemental Figure S3, B). These data suggest that R9 delivery is largely independent of cellular activity such as endocytosis. Co-infiltration of R9-GFP11 and endocytosis inhibitors wortmannin or ikarugamycin (Bandmann and Homann 2012, Bandmann, Müller et al. 2012, Elkin, Oswald et al. 2016) similarly resulted in no statistically significant decrease in delivery efficiency (Figure 3, F). Taken together, these data align with previously reported studies in mammalian cells and a singular plant protoplast centered study (Chugh and Eudes 2007) that suggest at concentrations greater than 10 μM, R9 enter cells through a combination of direct membrane permeation and through endocytosis and subsequent endosomal escape (Kosuge, Takeuchi et al. 2008, Wallbrecher, Ackels et al. 2017).

### The stability of peptide cargoes in leaves probed using DCIP

We next confirmed that DCIP is capable of detecting successful delivery in attached, intact leaves instead of leaf discs. DCIP expressing leaves were thus infiltrated with 100 μM R9-GFP11 solutions and allowed to incubate *in situ*. During this time, the infiltrated liquid would dry, rendering the effective treatment concentration difficult to compare to above leaf disc assays. We then assessed the amount of sfGFP complementation at 4H or 24H post infiltration with 100 μM R9-GFP11 (Figure 4, A). In accordance to the previous leaf disc assays, we observed relatively high percentage of sfGFP positive cells at 4H post infiltration in intact leaves. Conversely, at 24H post infiltration, the percentage of sfGFP positive cells returned nearly to baseline and the percent of GFP positive cells was not statistically different than the percent positive in the non-treated control. We also tested whether or not 24H treated cells were still competent to R9-mediated delivery; we retreated a previously 24H treated leaf with 100 μM R9-GFP11, and imaged 4H post re-treatment. In this re-treatment, we visually identified some instances of successful delivery (Figure 4, B). These data show that the signal observed from GFP11 delivery is transient but can be somewhat recovered with re-treatment.

**Figure 4.**
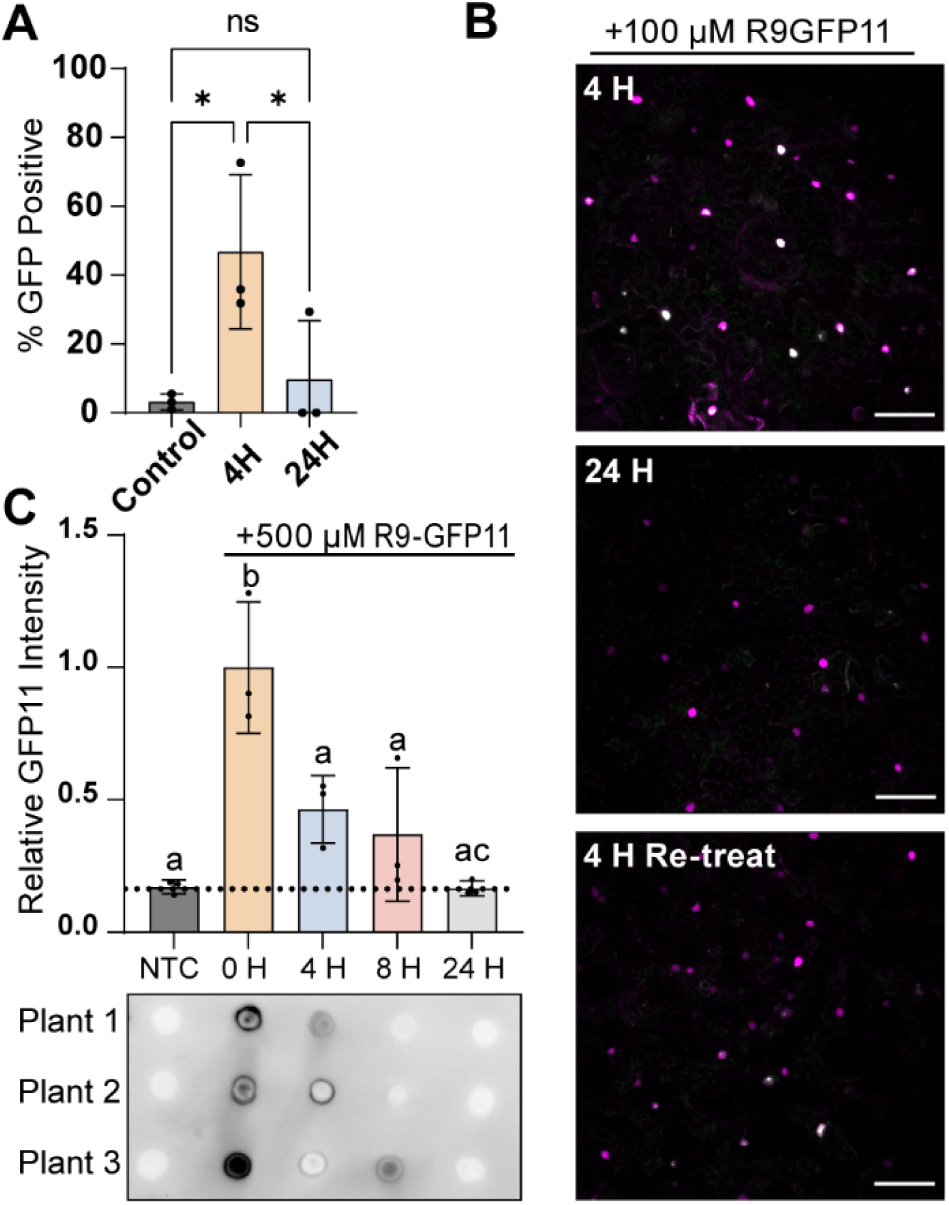
DCIP produces a transient response likely due to R9-GFP11 stability. A, Average percentage of GFP positive cells after infiltrating leaves with 100 µM R9-GFP11 for 4 or 24 Hours (N=3). Kruskal-Wallis test followed by Dunn’s multiple comparisons test was performed for all statistical comparisons where ns = p>0.05, * = 0.01<p<0.05. B, Representative two-color confocal maximum intensity projection micrographs of intact leaves treated with 100 µM R9-GFP11 at 4 H, 24 H, or retreated for 4 H post 24 H treatment. mCherry nuclei are pseudocolored magenta, and GFP green overlay results in white appearance. Scale bar is 100 µm. C, Dot blot of lysates containing GFP11 recovered from leaves infiltrated with 500µM R9-GFP11 or non-treated control for 0, 4, 8, and 24H and immunoblotted with an anti-GFP11 antibody. GFP11 spot intensity of non-treated leaves and leaves infiltrated with R9-GFP11 at 0, 4, 8, or 24H were normalized to the mean intensity of the 0H treatment to account of the variation in GFP11 recovery after lysis. Three separate plants were infiltrated for three biological replicates (N=3). Statistical comparison was performed with Kruskal-Wallis test where each letter represents groupings of statistical insignificance (p>0.05).

After DCIP revealed the transient delivery response, we sought to probe whether the stability of the R9-GFP11 construct contributed to the rapid peak and decline in response. We infiltrated leaves with 500 μM R9-GFP11 and left the plants to incubate *in situ*. A higher concentration of R9-GFP11 was chosen to facilitate detection by immunoblotting. After 0, 4, 8, and 24H of incubation, a single 12mm leaf disc was harvested and lysed for each treatment. Two microliters of lysates were then spotted onto nitrocellulose for dot blot analysis (Heinicke, Kumar et al. 1992) using a primary antibody raised against GFP11. The dot blot analysis shows rapid instability of GFP11 peptides in leaf tissues, with the quantity of recovered R9-GFP11 returning to levels of the non-treated control by 24H (Figure 4, C). These data suggest that the observed disappearance of DCIP signal is connected to clearance of the delivered peptide. Were the peptide stable in the tissue, we anticipate a continual delivery of the excess R9-GFP11 peptide from apoplast localized peptide. The reduction at 24H in sfGFP signal is indicative of the intracellular turnover of DCIP becoming higher than the rate of delivery. These data are in line with the limited serum stability of R9-conjugates in mammalian systems in which the limited peptide stability was associated with extracellular proteases and intracellular clearance of the peptide-cargoes (Palm, Jayamanne et al. 2007, Youngblood, Hatlevig et al. 2007). We hypothesize that similar dynamics from apoplastic proteases (Wang, Wang et al. 2020) and intracellular degradation are at play in plant cells. We would also add the additional caveat that the DCIP data may also be partially due to the intracellular half-life (∼26H) of GFP (Corish and Tyler-Smith 1999).

### DCIP enables qualitative detection of successful delivery of recombinant proteins

While the above quantification of CPP delivered peptides is useful, the delivery of larger constructs is required to facilitate the major goals of plant delivery. This motivated our design of a ligation independent cloning (Aslanidis and De Jong 1990) based *E. coli* expression vector for purifying recombinant proteins tagged with an N-terminal GFP11 and a C-terminal R9 peptide (Figure 5, A). To test the vector, we used mCherry (26.6 kDa) as a model protein. Although mCherry fluorescence overlaps with DCIP and precludes quantitative imaging, the identification of sfGFP fluorescent nuclei would confirm successful delivery. Furthermore, the innate mCherry fluorescence offers a glimpse of how well distributed the infiltrated protein solution is in the leaf tissue. We also purified an additional mCherry fusion cloned with a 3’ stop codon that would prevent tagging with the C-terminal R9 as a control. The total construct molecular weights were 32kDa without R9 and 34.5 kDa with R9.

**Figure 5.**
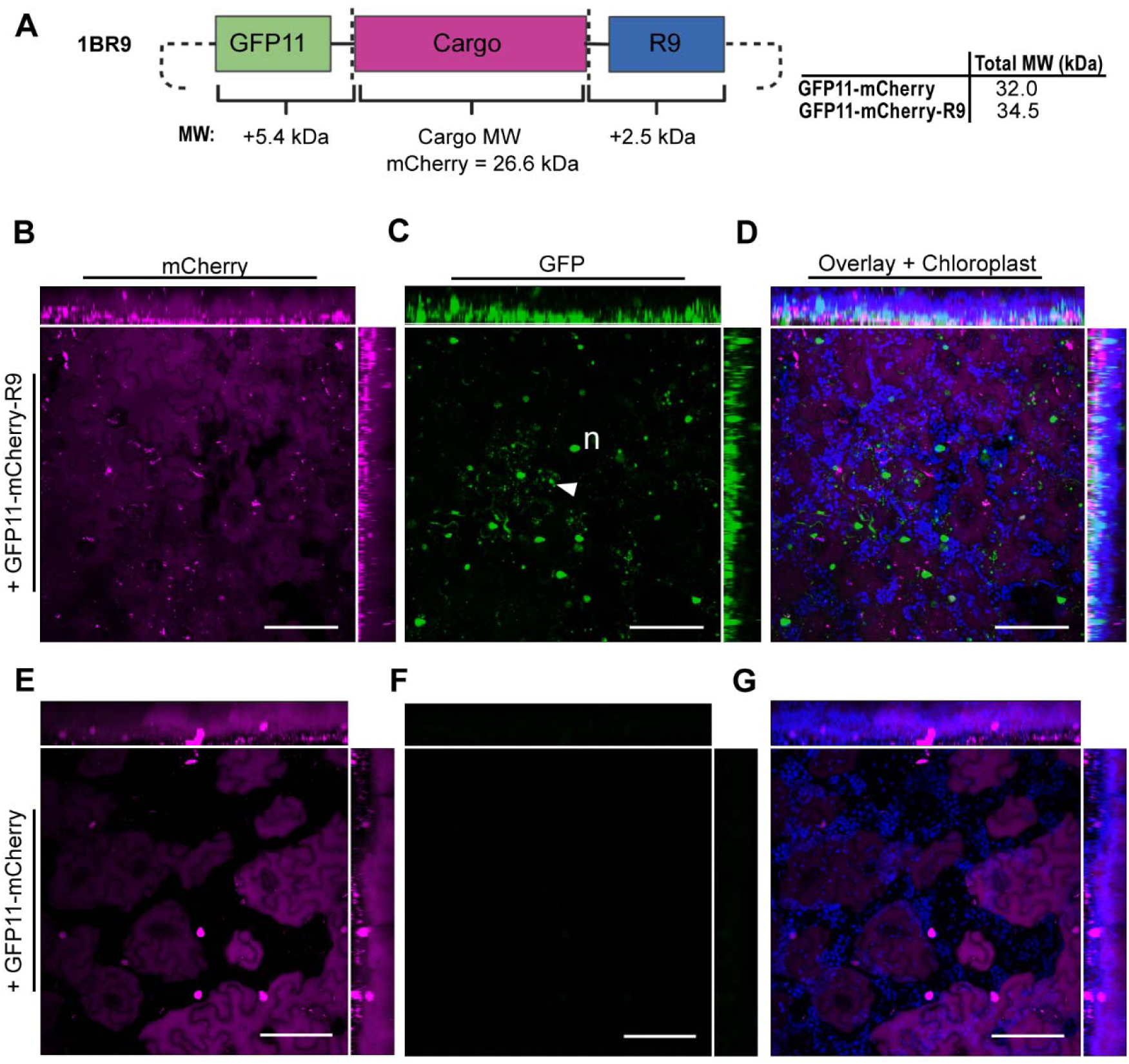
Qualitative confirmation of recombinant protein delivery. A, Schematic of LIC vector designed for tagging proteins of interest with N-terminal GFP11 and C-terminal R9. The resulting reporter and delivery tags result in an additional 7.9 kDa in MW to a recombinant protein of interest. B-D, DCIP sensor expressing plant infiltrated with 60µM GFP11-mCherry-R9 and incubated for 5H. B, mCherry fluorescence is pseudocolored magenta and demonstrates ubiquitous presence of infiltrated GFP11-mCherry-R9. C, sfGFP fluorescence from successful delivery is shown in green. Labeled features include nuclei (lower case “n”) and sfGFP containing intracellular aggregates or vesicular bodies (white triangle) resulting from successful delivery. D, An overlay of mCherry and GFP channels as well as chloroplast autofluorescence (pseudocolored blue). E-F, Equivalent experiment performed with 60µM GFP11-mCherry showing, E, mCherry fluorescence in magenta with some cells demonstrating strong DCIP nuclear fluorescence, F, sfGFP fluorescence (green), and, G, three color overlay with chloroplast channel. All images presented are maximum intensity projections with orthogonal projections of images. Scale bar is 100 µm.

As can be seen in Figure 5b & e, infiltration with 60µM of either mCherry construct results in thorough coverage of mCherry fluorescence. In GFP11-mCherry-R9 imaging, we were unable to discern the nuclear localized mCherry in DCIP from the infiltrated mCherry due to the bright signal of the infiltrated protein (Figure 5, B); although clear DCIP expression can be seen in the GFP11-mCherry treatment (Figure 5, E). This infiltration experiment also demonstrates the subjectivity of determining internalization from tissue-dispersed cargo fluorescence as the mCherry channel shows little obvious difference in appearance with or without R9 (Figure 5, B and E). Only with careful inspection can delivery be confirmed in the mCherry-R9 case by observing mCherry excluded zones caused by plastids (Supplemental Figure S8).

However, using the sfGFP channel, bright green fluorescent nuclei can be observed exclusively in the GFP11-mCherry-R9 infiltration (Figure 5, C). In addition to nuclei, we also observed numerous punctate sfGFP fluorescent objects that we hypothesize could be aggregates or partially entrapped GFP11-mCherry-R9 that have successfully undergone bimolecular fluorescence complementation. The observation of partial endosomal entrapment of mCherry aligns with the observation in mammalian cells that larger R9 delivered cargoes may undergo endocytosis and entrapment (Patel, Sayers et al. 2019). In contrast, without R9, no sfGFP fluorescence was observed (Figure 5, F), further confirming successful delivery of mCherry only when tagged with R9. An overlay of all channels with a chloroplast channel shows thorough delivery throughout the leaf tissue and that the green fluorescence does not arise from plastid autofluorescence (Figure 5, D and G). We also observed that successful delivery of the larger mCherry cargo appears qualitatively less efficient than delivery of smaller peptides (Supplemental Figure S9).

### Delivery of f-actin binding peptide with DCIP mediates protein-protein interactions in plants

After confirming that recombinant proteins could be efficiently delivered into plant cells with R9 CPP, we asked if delivered proteins could subsequently mediate protein-protein interactions. Being able to mediate protein-protein interactions using delivered proteins could enable new technologies that alter plant physiology. Therefore, in a proof-of-concept experiment, we delivered the 1.9kDa f-actin binding peptide, Lifeact (Riedl, Crevenna et al. 2008), tagged with GFP11 and R9 (Figure 6, A) into cytoDCIP expressing *N. benthamiana* leaves. In this scheme, when Lifeact is internalized, GFP11 acts as a scaffold to tether cytoDCIP to actin filaments through the Lifeact/f-Actin interaction. Kamiyama *et al* had previously shown that GFP11-sfGFP1-10 interactions could be used to mediate protein scaffolding in mammalian cells (Kamiyama, Sekine et al. 2016). Using our DCIP approach, we unambiguously confirm through imaging, that ectopic, delivery-mediated protein-protein interactions can be formed in plants. Previous studies using fluorescein labeled BP100-Lifeact had shown that delivered dye-labeled Lifeact could bind to plant actin in BY-2 cells. However, the constructs were not tested in leaves and did not leverage the low background and scaffolding afforded by a bimolecular fluorescence complementation approach (Eggenberger, Mink et al. 2011).

**Figure 6.**
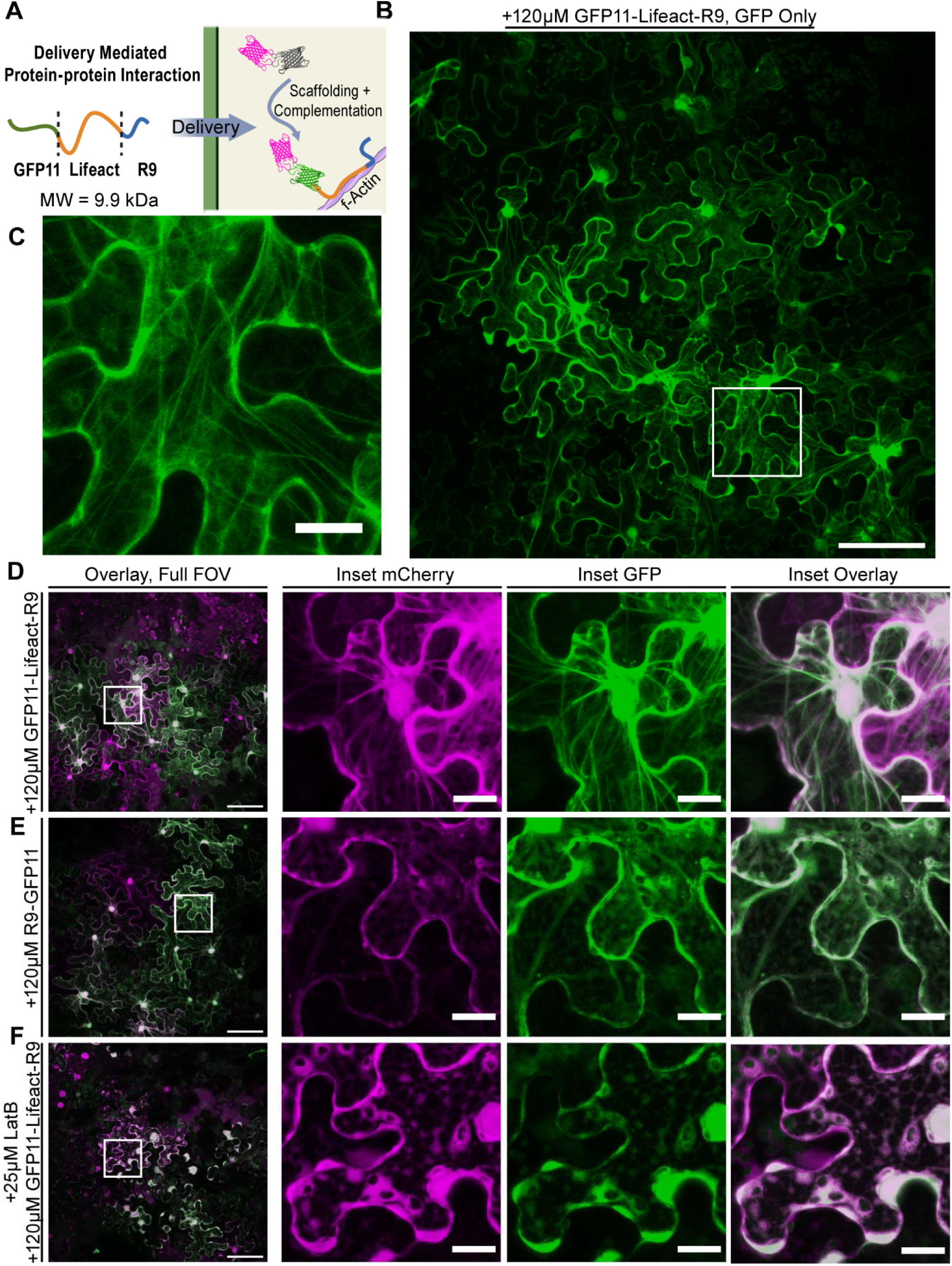
F-actin delivery-mediated protein-protein interactions formed in plants. A, Schematic of delivery mediated protein-protein interactions using purified recombinantly expressed GFP11-Lifeact-R9 (9.9 kDa). Infiltrated GFP11-Lifeact-R9 enters the cytosol through CPP-mediated delivery, binds to f-actin filaments in the plant cell, and scaffolds mCherry-sfGFP1-10 to the actin filament, thus mediating protein-protein interactions. B, Representative standard deviation projection FOV of actin labeling enabled by cytoDCIP and delivered GFP11-Lifeact-R9. CytoDCIP expressing leaves were infiltrated with 120 µM GFP11-Lifeact-R9 and incubated for 6H as leaf discs. C, Inset (white square) showing fine filament detail. D, Filaments appear as sfGFP fluorescent structured strands that colocalize with mCherry as result of cytoDCIP scaffolding. E, control treatment of CytoDCIP expressing leaves using 120 µM R9-GFP11 shows diffuse cytosolic localization of both mCherry and sfGFP. F, GFP11-Lifeact-R9 cotreated with 25µM Latrunculin B results actin depolymerization in diffuse cytosolic staining only. In all images, mCherry is pseudocolored magenta and sfGFP is pseudocolored green. Full FOV scale bar is 100 µm and inset 20 µm.

After a 6H incubation with infiltrated 120 µM GFP11-Lifeact-R9, we observe robust labeling of actin filaments (Figure 6, B) in pavement cells. At higher zoom, numerous fine structures are observed as the result of Lifeact delivery (Figure 6, C). Also importantly, we see colocalization of the mCherry signal with the sfGFP signal in the filamentous structures (Figure 6, D), suggesting that the cytoDCIP has been scaffolded to the actin filaments. When cytoDCIP expressing plants are treated with the molar equivalent of R9-GFP11, we observe no such fine structures and instead see diffuse, reticulated cytosolic localization. To further confirm that the observed filaments are indeed f-actin, we cotreated using either GFP11-Lifeact-R9 or R9-GFP11 with the f-actin destabilizing drug, latrunculin B (LatB) (Baluška, Jasik et al. 2001, Wright, Wood et al. 2007). In the presence of 25µM LatB, no actin filaments were observed (Figure 6, E), which suggests depolymerization of f-actin and confirms that the structures imaged were indeed the result of mCherry-sfGFP1-10 scaffolding to f-actin. We did however observe that LatB treatment changed the morphology of the cytosol in both the GFP11-Lifeact-R9 (Figure 6, E) treatment and the R9-GFP11 (Supplemental Figure S10). In both cases, the cytosol became granulated in pattern with the appearance of numerous, cytosol-excluding compartments. All together, these data suggest that delivered proteins could be used to perturb protein localization, mediate protein-protein interactions *in planta* and that DCIP or cytoDCIP may help accelerate identification of useful delivery mediated protein-protein interactions.

### Recombinant WUSCHEL delivery assisted by DCIP analysis

After establishing the feasibility of using DCIP to assess recombinant protein delivery in intact leaves, we sought to examine R9-mediated delivery of the *Arabidopsis* plant morphogenic transcription factor, WUSCHEL (AtWUS), in *N. benthamiana* leaves. AtWUS was chosen as a candidate cargo due to its applications for somatic embryogenesis in plants and its high degree of molecular characterization (Zuo, Niu et al. 2002, Ikeda, Mitsuda et al. 2009). DCIP expressing leaves infiltrated with 140 µM GFP11-AtWUS-R9 (MW = 41 kDa) showed robust nuclear GFP complementation at 6H (Figure 7, A). Because AtWUS lacks intrinsic fluorescence unlike mCherry, we were also able to demonstrate quantitative DCIP analysis using GFP11-AtWUS-R9 (Figure 7, B) with about 33% of DCIP expressing cells being GFP positive, suggesting a 33% AtWUS delivery efficiency on a per-cell basis. Because AtWUS is a transcription factor, we also expected to see GFP11-AtWUS-R9 to localize to the nucleus without the SV40 NLS of DCIP. To test this, we infiltrated a cytoDCIP expressing *N. benthamiana* leaf with 140µM GFP11-AtWUS-R9. If the delivered AtWUS has active NLS activity, we would expect green fluorescence localized to only the nucleus and excess, uncomplemented mCherry-GFP1-10 to remain in the cytosol. Indeed, at 6H we observe numerous GFP positive nuclei surrounded by cytosolic mCherry fluorescence (Figure 7, C), thus confirming native NLS targeting of delivered AtWUS. In contrast, R9-GFP11 treated cytoDCIP shows a general, cytosolic localization (Figure 6, E). These data show that R9 fusion is effective for WUS delivery and that the purified R9-tagged transcription factor is able to enter the nucleus.

**Figure 7.**
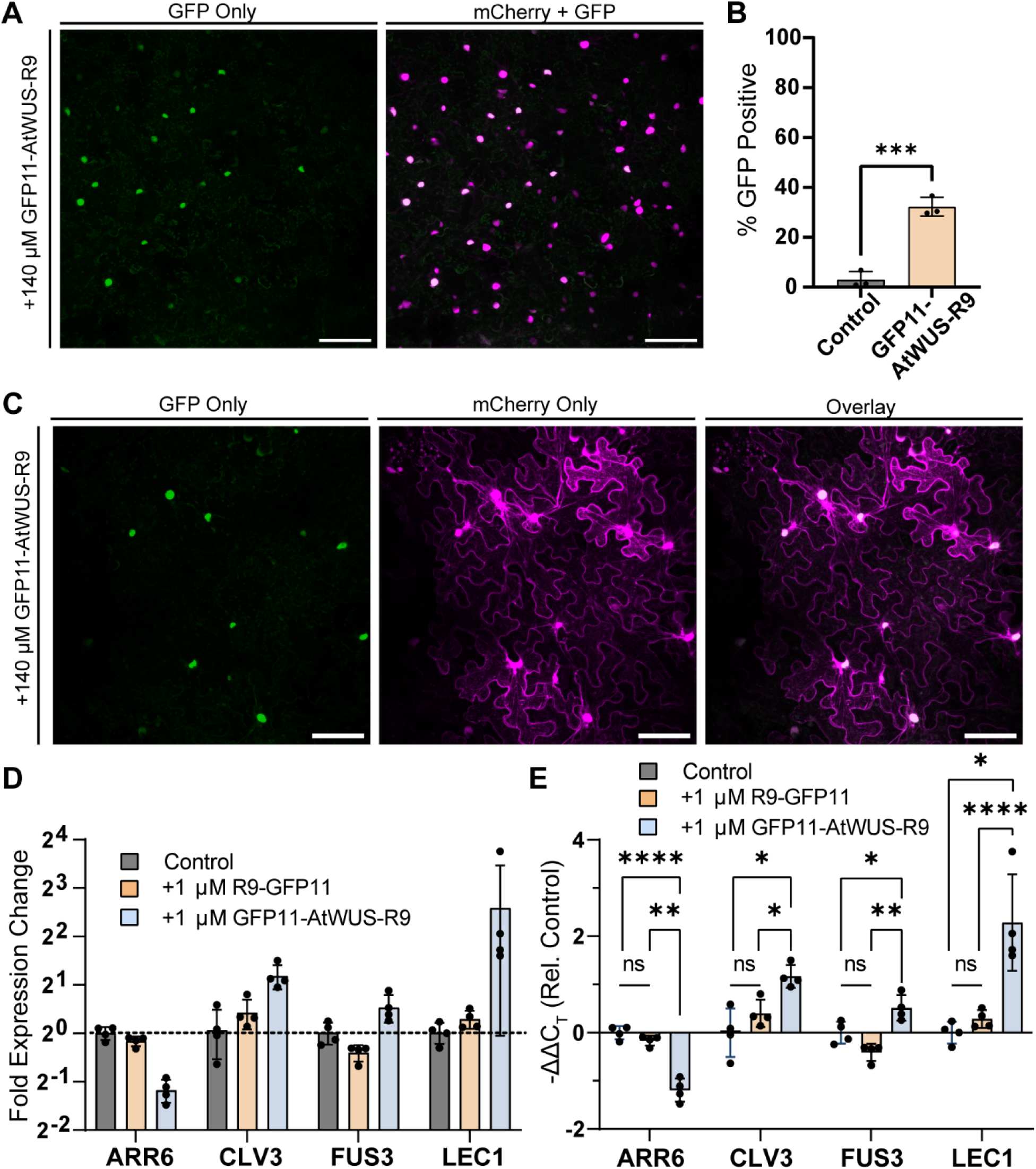
Delivery of the morphogenic transcription factor WUSCHEL. A, representative maximum intensity projection of a DCIP expressing *N. benthamiana* leaf infiltrated with 140µM GFP11-AtWUS-R9 and incubated as a leaf disc for 6H. Successful delivery presents as sfGFP (green pseudocolor) and mCherry (magenta pseudocolor) fluorescent nuclei as the result of delivered complementation. B, quantification of GFP positive nuclei as a result of AtWUS delivery in three plants (N=3) or buffer control. C, representative maximum intensity projection of cytoDCIP expressing *N. benthamiana* leaf infiltrated with 140µM GFP11-AtWUS-R9 and incubated as a leaf disc for 6H. The exclusive localization of sfGFP fluorescence and ubiquitous localization of cytosol localized cytoDCIP mCherry fluorescence shows that delivered GFP11-AtWUS-R9 is able to undergo native nuclear localization. D, rt-qPCR analysis of downstream AtWUS genes in 12-day old *Arabidopsis* seedlings treated with 1 µM GFP11-AtWUS-R9 or R9-GFP11 for 24H. 8-10 seedlings were treated per well and four wells were utilized for each treatment (N=4). E, statistical comparison showing measured ΔΔC_T_ values of GFP11-AtWUS-R9 treated seedlings are significantly changed when compared to either R9-GFP11 treatment or no treatment. Statistical analysis performed with T-test comparison and Holm-Šídák correction for multiple comparisons where ns = p>0.05, * = 0.01<p<0.05, ** = p<0.01, **** = p<0.0001.

The previous results in *N. benthamiana* motivated us to determine whether or not the delivered AtWUS is transcriptionally active. We treated 12-day-old *Arabidopsis* seedlings for 24H with 1µM GFP11-AtWUS-R9. *A. thaliana* was chosen as a model species due to the well characterized AtWUS pathway in *A. thaliana*. Seedlings were subsequently harvested and subjected to RT-qPCR analysis for several known downstream targets of AtWUS (Figure 7, D). GFP11-AtWUS-R9 delivery led to down-regulation of ARR6 (0.44-fold) and upregulation of CLV3 (2.26-fold), FUS3 (1.44-fold), and LEC1 (6-fold) when compared to the buffer-treated control. Statistical analysis of the measured C_T_ values show that R9-GFP11 alone does not mediate significant transcriptional changes while GFP11-AtWUS-R9 recapitulates the expected transcriptional response to AtWUS overexpression (Figure 7, E). Ectopic expression of AtWUS downregulates the expression of ARR6 (Leibfried, To et al. 2005), and upregulates the expression of its own negative regulator, CLV3 (Schoof, Lenhard et al. 2000). Additionally, AtWUS overexpression also upregulates transcripts of somatic embryogenesis markers, FUS3 and LEC1 (Ikeda, Mitsuda et al. 2009). These data suggest that not only is AtWUS delivery possible, but that this delivered transcription factor can be transcriptionally active in plants.

## Discussion

Protein delivery in plants is an emerging field motivated by the need for better handles to dissect plant molecular physiology, enable DNA-free gene editing technology, and enhance agronomic traits. Previous foundational work by Numata *et al* have identified cell penetrating peptides as a method to deliver proteins using a dye-mediated approach (Numata, Horii et al. 2018). However, the journey from the apoplast into the cytoplasm causes cargoes to be excluded, entrained, or sequestered and necessitates a cargo-in-cytosol dependent approach. Furthermore, technical limitations render microscopic analysis of uptake only *suggestive* of uptake rather than *confirmative*, and preclude quantification of relative CPP delivery efficiencies. Previously, final confirmation of delivery in plants has required lengthy functional analysis such as identifying delivery mediated gene editing (Martin-Ortigosa, Peterson et al. 2013), protein expression (Demirer, Zhang et al. 2018, Thagun, Horii et al. 2022), or silencing (Zhang, Goh et al. 2022). We therefore sought to develop a technique by which delivery designs could be rapidly and accurately assessed, shown here for CCP-mediated peptide and protein delivery, but could be generically extended to testing of other carriers. Although a suite of tools has been developed for use in mammalian cell culture to confirm CPP-mediated cytosolic delivery (Schmidt, Adjobo-Hermans et al. 2015, Peraro, Deprey et al. 2018, Teo, Rennick et al. 2021), no such tool has been tailored for plants until the present work.

Our development, delivered complementation *in planta* (DCIP) is a microscopic tool that unambiguously confirms the delivery of peptides and proteins even at concentrations as low as 10 µM in leaves. DCIP also enables quantitative measurement of relative delivery efficiency, thus enabling a new functional method to rapidly screen effective cell penetrating peptides. In our assay, we identified TAT and R9 as being effective *in planta* delivery CPPs, with R9 as the most effective, whereas the previously identified BP100 (Numata, Horii et al. 2018) was ineffective at delivering GFP11. One potential explanation for this discrepancy is the sensitivity of CPPs to chemical conjugation and that dye-conjugation as used in previous CPP screens perturbs their uptake performance (Birch, Christensen et al. 2017). The use of GFP11 in DCIP as a reporter tag presents an improvement to screening strategies as it provides a clear, low-background signal of successful cytosolic delivery. We additionally used DCIP to show that uptake of R9 in plants is endocytosis-independent in line with previous literature in mammalian systems (Brock 2014). Unraveling the CPP uptake mechanism in plants may lead to novel strategies for improving peptide and protein delivery *in planta*. These experiments show DCIP is a tool that can be used both to interrogate delivery and also to screen novel CPP sequences and chemistries that could improve delivery of not just proteins but eventually RNA or DNA.

DCIP has also enabled us to study the durability of delivered peptides *in planta*. We used DCIP and observed that the sfGFP complementation response from R9-GFP11 delivery was transient and disappeared by 24H. We orthogonally confirmed that R9-GFP 11 is cleared from leaf tissue by 24H with immunoblotting. These results suggest that the stability of cargo will be an important consideration for designing effective plant bioengineering strategies that leverage delivery, and that successful delivery is also contingent on cargo stability protection in addition to delivery itself. These results align with previous nanoparticle mediated delivery strategies that involve stabilization of the cargo to lytic enzymes but not necessarily cellular internalization of the vehicle itself (Zhang, Goh et al. 2022). DCIP therefore offers a facile tool for engineering novel carrier-based designs that optimize both delivery efficiency and cargo stability.

Next, we used DCIP and a cytosolically localized variant, cytoDCIP, to investigate the delivery of larger recombinant proteins. We show that GFP11-mCherry can be delivered using a c-terminally fused R9 CPP to leaf cells. In contrast to previous recombinant fluorescent protein delivery strategies, the bimolecular fluorescence complementation signal we observe can only occur upon successful cytosolic delivery and is not convolved with apoplastic or endosomally entrapped material. As a proof of concept that future experiments that leverage delivery-mediated protein-protein interactions may be possible, we then used cytoDCIP to show that specific protein-protein interactions can be mediated through a delivered recombinant protein by tethering the mCherry-sfGFP1-10 to f-actin through Lifeact/f-actin binding. Incidentally, the development of GFP11-Lifeact-R9 may also provide novel way to visualize actin filaments in plants.

As a final demonstration for the usefulness of DCIP, we show, to our knowledge, the first delivery of a plant transcription factor, AtWUS, to walled plant cells. We not only show, through imaging, that it is possible to deliver recombinant AtWUS to *N. benthamiana* but also show, through RT-qPCR, that delivered AtWUS recapitulates AtWUS overexpression transcriptional downstream responses in *Arabidopsis* seedlings. DNA-based overexpression of AtWUS and its orthologs has been found to enhance the regeneration of transgenics and somatic embryogenesis of numerous species such as cotton, sorghum and maize (Bouchabké-Coussa, Obellianne et al. 2013, Lowe, Wu et al. 2016, Che, Wu et al. 2022). The confirmation of active AtWUS delivery evinces a new DNA-free strategy for enhancing the recovery of transgenic plants without DNA-based WUS overexpression and subsequent transgene excision in challenging species. However, future work will be needed to develop a strategy for delivered WUSCHEL mediated somatic embryogenesis.

In summary, our development and validation of DCIP, **D**elivered **C**omplementation ***i****n **p**lanta*, could enable the design of novel cell penetrating peptides or nanoparticle-based carriers for protein delivery. For example, DCIP might be used to identify plant-specific cell penetrating motifs from secreted effector proteins of pathogenic fungi (Bouwmeester, Meijer et al. 2011). DCIP may also be used in future experiments to determine whether larger proteins such as Cas9 or novel miniaturized Cas12f variants could be delivered (Xu, Chemparathy et al. 2021). In addition to aforementioned DNA-free gene editing, DCIP could engender the delivery nanobodies for pathogen resistance and targeted protein degradation (Caussinus, Kanca et al. 2012, Hemmer, Djennane et al. 2018), the delivery of stress tolerance conferring disordered proteins (Wallmann and Kesten 2020), or the delivery of a greater variety of morphogenic regulators to control plant regeneration.

## Methods

### Reagents and Antibodies

Reagents, buffers, and media components were procured through Sigma-Aldrich unless otherwise noted. Solid-phase chemical peptide synthesis of GFP11 and CPP fusions was performed by a third-party manufacturer (GenScript). Enzymes used for cloning reactions were procured through New England Biolabs. Anti-GFP11 antibody was purchased through Thermo-Fisher and the anti-mCherry and anti-rabbit Igg-HRP secondary antibody through Cell Signaling Technologies. All oligonucleotides and DNA sequences were purchased from Integrated DNA Technologies (IDT).

### Plant growth conditions and agroinfiltration

*N. benthamiana* were grown in a growth chamber kept at 24 °C and a light intensity of 100-150 µmol_m^−2^_s^−1^. The photoperiod was kept at 16H light/8H dark. Seeds were sown in inundated soil (Sunshine Mix #4) and left to germinate for 7-10 days at 24°C before being transferred to 10cm pots for growth. Fertilization was done on a weekly basis with 75 ppm N 20-20-20 general-purpose fertilizer and 90 ppm N calcium nitrate fertilizer reconstituted in water. Infiltrations were performed on 4–5-week-old plants on the third and fourth expanded leaves. Agroinfiltrations were performed via needless syringe using overnight cultures of *A. tumefasciens* bearing the DCIP constructs. The day of infiltration, the overnight 30°C cultures were pelleted at 3200xg, rinsed with infiltration buffer (10mM MES pH 5.7, 10mM MgCl_2_) and then resuspended in infiltration buffer containing 200 μM acetosyringone to an OD of 0.5-1.0. The cultures were then left shaking at ambient conditions for 2-4 hours before final adjustment of the OD_600_ to 0.5 with infiltration buffer. During infiltration, care was taken to minimize the number of damaging infiltration spots to completely saturate the leaf. The plants were then left at ambient conditions overnight to dry before being transferred to the growth chamber for the total incubation time of 3 days.

### Plasmid Construction and Bacterial Strains

A list of parent plasmids and newly constructed plasmids, are included in Supplemental Table 1. Additionally, the predicted protein products and organization of all constructed plasmids are provided. Primers and synthetic DNA used for cloning are included in Supplemental Table 2. For all cloning steps, plasmids were transformed into XL1-blue *E. coli*. The coding sequence of DCIP was constructed by ligating mCherry sequence into a PstI 5’ of the sfGFP1-10 coding sequence in pPEP101 (Park, Lee et al. 2017). In the nuclear localized variant of DCIP, NLS was attached during PCR of the mCherry sequence. The NLS is omitted in cytoDCIP. Next, the coding sequences of DCIP and cytoDCIP were amplified by PCR for domestication into pUDP2 before final Golden Braid (GB2.0) assembly following the standard GB2.0 protocol using Esp3I (Sarrion-Perdigones, Vazquez-Vilar et al. 2013). The DCIP and cytoDCIP transcriptional units were assembled using pUDP2-35S-oTMV for the promoter and pUDP2-tNOS for the terminator in a BsaI restriction-ligation GB2.0 reaction. cytoDCIP and DCIP were then transformed into GV3101 *Agrobacterium tumefasciens* bearing pSOUP (Hellens, Edwards et al. 2000) and plated onto LB agar containing rifampicin (50μg/mL), gentamicin (25μg/mL), and kanamycin (50 μg/mL).

The recombinant GFP1-10 expression vector, 1B-GFP1-10, was constructed by PCR amplifying the sfGFP1-10 gene from a pPEP101 with the ligation-independent cloning tags for plasmid 1B. The full protocol for LIC cloning used was provided by the UC Berkeley Macro Lab: https://qb3.berkeley.edu/facility/qb3-macrolab/projects/lic-cloning-protocol/. The resulting amplicon was then inserted by LIC into 1B and transformed into *E. coli* for expansion, purification, sequencing, and transformation into an expression *E. coli* strain. The 1BR9 plasmid was constructed by inserting a short, chemically synthesized DNA sequence containing N-terminal GFP11 and C-terminal R9 tag into plasmid 1B via LIC. Between the tags, a new LIC site was regenerated such that future LIC reactions would insert the protein of interest between the N- and C-tags. The LIC approach allowed insertion of PCR amplified mCherry sequence into 1BR9 to generate 1BR9-mCherry. Incorporation of a TAA stop codon into the reverse primer generated 1BR9-mCherrySTOP which excludes the c-terminal R9 motif. 1BR9-Lifeact was produced by inserting a chemically synthesized DNA sequence for Lifeact into 1BR9. 1BR9-AtWUS was constructed by LIC insertion of an *E. coli* codon optimized (IDT) synthesized DNA AtWUS (TAIR: At2g17950.1) DNA sequence into the 1BR9 vector.

### Recombinant Protein Expression and Purification

For all recombinant protein expression, Rosetta 2 (DE3) pLysS *E. coli* were transformed with recombinant protein expression vectors and plated on to selective chloramphenicol (25 μg/mL) and kanamycin (50 μg/mL) LB agar plates for overnight growth at 37°C, 250rpm. Single colonies were used to inoculate 10mL seed cultures in LB. After overnight starter culture growth at 37°C and 250rpm, 1L LB with selective antibiotics were set to grow at 37°C, 250rpm in 2L baffled flasks. Induction was performed with 0.5 mM IPTG at 37°C when the culture reached 0.8 OD_600_. After 4 hours of induction, cultures were pelleted for 20 minutes at 3200xg and flash frozen in liquid nitrogen. Cell lysis was conducted using thawed pellets in lysis buffer (50mM Tris-HCL, 10mM imidazole, 500mM NaCl, pH 8.0) with 1x protease inhibitor cocktail (Sigma-Aldrich: S8830) using probe tip sonication. The resulting lysate was clarified by centrifugation at 40,000xg for 30 minutes. For GFP1-10 and mCherry fusion proteins, the soluble fraction was incubated with 1mL Ni-NTA (Thermo Scientific: 88221) slurry for an hour. After incubation and washing, the proteins were eluted using elution buffer (500mM imidazole, 150mM Tris-HCL, pH 8.0). The resulting eluate was then concentrated and buffer exchanged into 10mM Tris-HCL, 100mM NaCl pH 7.4 via ultrafiltration in a 3500Da cutoff filter (Emdmillipore: C7715). The GFP11-mCherry and GFP11-mCherry-R9 were then further polished using SEC (Cytiva: HiLoad 16/600 Superdex 200pg) and exchanged into storage buffer (10mM Tris, 10mM NaCl pH 7.4) before ultrafiltration concentration and flash freezing for storage. For GFP11-Lifeact-R9, the insoluble pellet from clarification was solubilized in 8M Urea, 50mM Tris, pH 8.0 before incubation with Ni-NTA for an hour. In addition to standard wash steps, a high pH wash (20mM CAPS pH 10.8, 1M NaCl) was required to remove residual nucleic acids from the protein. Protein was then eluted with elution buffer before spin concentration and exchange into storage buffer and flash freezing. Aliquots of each recombinant protein were run on SDS-PAGE for confirmation (Supplemental Figure S11).

### Recombinant GFP11-AtWUS-R9 Purification

GFP11-AtWUS-R9 was expressed as mentioned previously. However, purification proceeded by sonication lysis in 6M Urea, 50mM Tris, 0.5mM TCEP, and 2mM MgCl_2_ pH 7.5 in the presence of 25 U/mL of benzonase and protease inhibitor cocktail. After lysis, the lysate was then incubated at 37C for 30 minutes with occasional mixing. After incubation, an additional 25 U/mL of benzonase was added and the lysate was clarified by centrifugation at room temperature, 40,000xg for 30 minutes. To the clarified supernatant, 2mL of Ni-NTA slurry per liter of starting culture was added and incubated at room temperature for 2 hours. The Ni-NTA was then washed sequentially with at least 15 bed volumes each of wash A (6M Urea, 50mM Tris pH 7.5), then wash B (6M Urea, 50mM Tris, pH 7.5, 500mM NaCl), then wash C (6M Urea, 50mM Tris, pH 7.5, 50mM Imidazole). The protein was eluted twice with the addition of 1 bed volume of 1M Imidazole, 20mM MES, pH 6.9, 200mM NaCl. The fractions were then combined and buffer exchanged into buffer P (30mM Sodium Acetate, 5mM NaCl, pH 4.0, 0.5mM TCEP) by dialysis or by desalting in a G25 sephadex pre-packed PD-10 column (Cytiva) and then analyzed with SDS-PAGE and *in vitro* complementation (Supplemental Figure S12).

### *In vitro* GFP complementation assay

In a 96-well qPCR plate (Biorad), 5 µL of 10 µM total protein of sfGFP1-10 Ni-NTA eluate in storage buffer (10mM Tris, 10mM NaCl pH7.4) was combined with 5µL buffer or 20 µM of each tested peptide or protein in storage buffer. Each well was mixed by pipetting and was tested in triplicate. GFP complementation of GFP11-AtWUS-R9 was accomplished similarly except 10µL of 15µM either GFP11-AtWUS-R9 or R9-GFP11 in buffer P was added to 10µL of 10µM GFP1-10. An additional 5µL 1M Tris pH 8.0 was also added to overcome the acidity of buffer P. GFP complementation was quantified using a Biorad CFX96 qPCR machine by measuring the green fluorescence at 1-minute intervals over the course of 6 hours at either 22°C or 4°C.

### SDS-PAGE and Western Blot

For Western Blot of DCIP and cytoDCIP, agroinfiltrated leaves were harvest 3 d.p.i, flash frozen, ground, and lysed with RIPA buffer (Abcam: ab156034) for 20 minutes. Lysates were then clarified by centrifugation and boiled in 1X Laemmli buffer (Biorad) and 10 (V/V) % beta-mercaptoethanol for 5 minutes before being loaded into a 4-20% gradient SDS-PAGE gel (Biorad: 4561096) and run according the manufacturer’s instructions. Membrane transfer was done according to manufacturer’s instructions onto Immobilon PVDF membrane (Millipore). Blocking was performed using 5% milk in PBS with 0.1% Tween (PBST). The incubation with primary anti-mCherry antibody (CST: E5D8F) was performed overnight at 1:1000 dilution in 3% BSA in PBST at 4°C with orbital shaking at 60 rpm. Imaging was performed after probing with anti-Rabbit IGG-HRP secondary antibody (CST: 7074) at 1:10,000 dilution in 5% milk PBST and ECL prime chemiluminescent reagent (Amersham: RPN2236) on a ChemiDoc gel imager (Biorad).

### R9-GFP11 Dot Blot Assay

The third or fourth leaf of 4–5-week-old wild-type *N. benthamiana* were infiltrated with 500 μM R9-GFP11 in water using a needless syringe. Infiltrations were staggered such that all treatments could be harvested simultaneously for 0, 4, 8, and 24H time points. Each treatment was performed on a separate plant and the experiment was repeated thrice. A 12mm leaf disc was excised for each treatment using a leaf punch and flash frozen in liquid nitrogen before grinding and lysis in 20 μL RIPA buffer with 1x plant protease inhibitor cocktail (Sigma-Aldrich: P9599). The lysates were then clarified by centrifugation at 21000xg for 30 minutes. Immediately after, 2 μL lysates were spotted onto a nitrocellulose membrane (Amersham: GE10600002) and allowed to dry. The membranes were then blocked in 5% milk in PBS-T before being washed and probed with an anti-GFP11 antibody (Invitrogen: PA5-109258) (1:500 dilution in PBST with 3% BSA). Secondary antibody probing and imaging was performed similarly to Western Blot.

### Delivered complementation *in planta* infiltration

Three days after agroinfiltration, DCIP expressing leaves were infiltrated with the treatment solutions. Unless otherwise stated, an 8mm punch was then excised from the infiltrated area and plated, abaxial side up, onto ½ MS pH 5.7 agar plates. The plates were then left to incubate under ambient conditions for 4-5 H before imaging. For cold temperature treatment, after infiltration with ice-cold solutions of R9-GFP11 and disc excision, the leaf discs were plated onto ice-cold agar and immediately transferred to a 4°C refrigerator for incubation. After incubation, the agar plates with leaf discs were kept on ice until the moment of imaging. For *in situ* incubation, leaves were simply infiltrated and the plant was returned to the growing chamber for incubation. All peptides were dissolved in sterile water and all recombinant proteins were dissolved in 10mM Tris pH 7.4, 10mM NaCl for infiltration. GFP11-AtWUS-R9 was found to possess low solubility at neutral pH and was exchanged into 10mM MES, pH 5.5, immediately before infiltration.

### Confocal Imaging and Image Analysis

Excised leaf discs were imaged on a Zeiss LSM880 laser scanning confocal microscope. Images for semi-quantitative analysis were collected using a 20x/1.0NA Plan-apochromat water immersion objective and larger field-of-views were collected using a 5x objective. Leaf discs were mounted by sandwiching a droplet of water between the leaf disc and a #1.5 cover glass. sfGFP, mCherry, and chloroplast autofluorescence images were acquired by excitation with a 488, 561, and 635 nm laser respectively. The emission bands collected for sfGFP, mCherry, and autofluorescence were 493-550 nm, 578-645 nm and 652-728 nm respectively. All images were collected such that the aperture was set to 1 Airy-unit in the sfGFP channel. Images were prepared for publication using Zen Blue software. For quantification experiments, z-stacks were acquired with the imaging depth set to capture the epidermal layer down to the point where mCherry nuclei could no longer be detected. Z-stacks from four field of views were acquired for every treatment condition. Quantitative image analysis and downstream processing was performed using Cell Profiler 3.0. On a slice-by-slice basis, mCherry fluorescent nuclei were segmented using Otsu’s method (Otsu 1979) and chloroplasts were segmented in the autofluorescence channel. The segmented chloroplasts were applied as a sfGFP channel mask over the image to exclude plastid autofluorescence in downstream image analysis. After masking, maximum intensity projections of identified nuclei were generated in the sfGFP, autofluorescence, and mCherry channels. Cell Profiler was then used to quantify the number, mCherry intensity, GFP intensity, and red (mCherry)/green (sfGFP) ratio of the projected nuclei.

### Arabidopsis GFP11-AtWUS-R9 Seedling Treatment

Arabidopsis (Col-0) seedlings were grown in 12-well culture plates with 8 to 10 seedlings per well in 1 mL of 1x MS media supplemented with 0.5% sucrose and 2.5 mM MES, pH 5.7. Seeds were sterilized by washing in 70% ethanol for 30 seconds followed by a 15-minute incubation in 50% bleach supplemented with 0.5% Tween-20 and rinsed 5x with DI water. Sterilized seeds were stratified in plates at 4°C for 3 days after plating. Seedlings were grown at 22°C under 16-hour photoperiods for 12 days. To treat seedlings, the liquid media in each well was replaced with control and Wuschel treatments. Control wells were refreshed with 1 mL of MS growth media. Treatment wells were refreshed with 1 mL of 1 μM of protein (R9-GFP11 or R9-WUS-GFP11) dissolved in MS growth media. After 24 hours of treatment, seedlings were frozen in liquid nitrogen and physically disrupted with chrome steel bearing balls.

### RT-qPCR analysis

Total RNA was extracted from Arabidopsis seedlings using a RNeasy Plant Mini Kit (Qiagen). Extracted RNA quality was confirmed with a NanoDrop UV-Vis Spectrometer. Complementary DNA was synthesized from RNA using an iScript cDNA Synthesis Kit (Bio-Rad). qPCR was run with PowerUP SYBR Green Master Mix (Applied Biosystems), each reaction was run in triplicate according to manufacturer’s recommendations. Melt-curve analysis was run after qPCR cycling to confirm primer specificity. Relative gene expression was determined using the ddCt method (Livak and Schmittgen 2001) using SAND1 as the reference gene. Relative gene expression was determined from 4 biological pools each containing 8 to 10 seedlings. A list of utilized primers is available in (Supplemental Table 2). Statistical comparisons of ddC_T_ values were conducted as done previously (Yuan, Reed et al. 2006) using a T-test with Holm-Šídák correction in GraphPad Prism 9.

### Data Processing and Statistics

Data obtained from Cell Profiler was processed using a script written in Python 3.9. Using the Cell Profiler data, GFP positive nuclei were counted using a python script by setting a threshold defined as a one-tailed 99% confidence interval above the mean green/red ratio in the untreated control or water infiltration control of each experiment. Any nuclei with green/red ratio higher than this threshold would be identified as GFP positive. The percentage of sfGFP positive nuclei was calculated by dividing sfGFP positive nuclei by the total number of mCherry nuclei counted. In experiments where delivery efficiency is used, delivery efficiency is defined by normalizing the percentage of sfGFP positive nuclei to the 100μM R9-GFP11 treatment in that experiment. All summary statistics were calculated using Python before export and statistical analysis in GraphPad Prism 9. Kruskal-Wallis non-parametric ANOVA (Kruskal and Wallis 1952) was used for analysis of multiple comparisons followed by uncorrected Dunn’s non-parametric T-test. Single comparisons were made using a one-sample t-test against the normalized value of 1.0. All presented plots were also generated in GraphPad Prism 9.

## Acknowledgements

We would like to thank the lab of Prof. Savithramma Dinesh-Kumar for provisioning the initial plasmids for sfGFP1-10. We thank the UC Berkeley Macrolab for providing the backbone vector 1B and LIC cloning protocol. We also would like to thank Antonio Del Rio Flores for advice and support for protein purification. J.W.W. is supported by NSF GRFP. N.S.G. is supported by a FFAR Fellowship.. Confocal imaging experiments were conducted at the CRL Molecular Imaging Center, RRID:SCR_017852, supported by the Helen Wills Neuroscience Institute. We would like to thank Holly Aaron and Feather Ives for their microscopy advice and support. We further acknowledge support of a Burroughs Wellcome Fund Career Award at the Scientific Interface (CASI) (to M.P.L.), a Dreyfus foundation award (to M.P.L.), the Philomathia foundation (to M.P.L.), an NIH MIRA award (to M.P.L.), an NIH R03 award (to M.P.L), an NSF CAREER award (to M.P.L), an NSF CBET award (to M.P.L.), an NSF CGEM award (to M.P.L.), a CZI imaging award (to M.P.L), a Sloan Foundation Award (to M.P.L.), a USDA BBT EAGER award (to M.P.L), a Moore Foundation Award (to M.P.L.), an NSF CAREER Award (to M.P.L), and a DOE office of Science grant with award number DE-SC0020366 (to M.P.L.). M.P.L. is a Chan Zuckerberg Biohub investigator, a Hellen Wills Neuroscience Institute Investigator, and an IGI Investigator.

## References

Aslanidis, C. and P. J. De Jong (1990). “Ligation-independent cloning of PCR products (LIC-PCR).” Nucleic acids research 18(20): 6069–6074.

Baluška, F., J. Jasik, H. G. Edelmann, T. Salajová and D. Volkmann (2001). “Latrunculin B-Induced Plant Dwarfism: Plant Cell Elongation Is F-Actin-Dependent.” Developmental Biology 231(1): 113–124.

Bandmann, V. and U. Homann (2012). “Clathrin-independent endocytosis contributes to uptake of glucose into BY-2 protoplasts.” The Plant Journal 70(4): 578–584.

Bandmann, V., J. D. Müller, T. Köhler and U. Homann (2012). “Uptake of fluorescent nano beads into BY2-cells involves clathrin-dependent and clathrin-independent endocytosis.” FEBS Letters 586(20): 3626–3632.

Birch, D., M. V. Christensen, D. Staerk, H. Franzyk and H. M. Nielsen (2017). “Fluorophore labeling of a cell-penetrating peptide induces differential effects on its cellular distribution and affects cell viability.” Biochimica et Biophysica Acta (BBA) - Biomembranes 1859(12): 2483–2494.

Bouchabké-Coussa, O., M. Obellianne, D. Linderme, E. Montes, A. Maia-Grondard, F. Vilaine and C. Pannetier (2013). “Wuschel overexpression promotes somatic embryogenesis and induces organogenesis in cotton (Gossypium hirsutum L.) tissues cultured in vitro.” Plant Cell Reports 32(5): 675–686.

Bouwmeester, K., H. J. G. Meijer and F. Govers (2011). “At the Frontier; RXLR Effectors Crossing the Phytophthora–Host Interface.” Frontiers in Plant Science 2.

Brock, R. (2014). “The uptake of arginine-rich cell-penetrating peptides: putting the puzzle together.” Bioconjugate chemistry 25(5): 863–868.

Caussinus, E., O. Kanca and M. Affolter (2012). “Fluorescent fusion protein knockout mediated by anti-GFP nanobody.” Nature Structural & Molecular Biology 19(1): 117–121.

Chang, M., J.-C. Chou, C.-P. Chen, B. R. Liu and H.-J. Lee (2007). “Noncovalent protein transduction in plant cells by macropinocytosis.” New Phytologist 174(1): 46–56.

Che, P., E. Wu, M. K. Simon, A. Anand, K. Lowe, H. Gao, A. L. Sigmund, M. Yang, M. C. Albertsen, W. Gordon-Kamm and T. J. Jones (2022). “Wuschel2 enables highly efficient CRISPR/Cas-targeted genome editing during rapid de novo shoot regeneration in sorghum.” Communications Biology 5(1): 344.

Chugh, A. and F. Eudes (2007). “Translocation and nuclear accumulation of monomer and dimer of HIV-1 Tat basic domain in triticale mesophyll protoplasts.” Biochimica et Biophysica Acta (BBA) - Biomembranes 1768(3): 419–426.

Corish, P. and C. Tyler-Smith (1999). “Attenuation of green fluorescent protein half-life in mammalian cells.” Protein Engineering, Design and Selection 12(12): 1035–1040.

Demirer, G. S., H. Zhang, N. S. Goh, R. L. Pinals, R. Chang and M. P. Landry (2020). “Carbon nanocarriers deliver siRNA to intact plant cells for efficient gene knockdown.” Science Advances 6(26): eaaz0495.

Demirer, G. S., H. Zhang, J. Matos, N. Goh, F. J. Cunningham, Y. Sung, R. Chang, A. J. Aditham, L. Chio, M.-J. Cho, B. Staskawicz and M. P. Landry (2018). “High Aspect Ratio Nanomaterials Enable Delivery of Functional Genetic Material Without DNA Integration in Mature Plants.”

Dolde, U., V. Rodrigues, D. Straub, K. K. Bhati, S. Choi, S. W. Yang and S. Wenkel (2018). “Synthetic MicroProteins: Versatile Tools for Posttranslational Regulation of Target Proteins.” Plant Physiology 176(4): 3136–3145.

Donaldson, L. (2020). “Autofluorescence in Plants.” Molecules (Basel, Switzerland) 25(10): 2393.

Eggenberger, K., C. Mink, P. Wadhwani, A. S. Ulrich and P. Nick (2011). “Using the peptide bp100 as a cell-penetrating tool for the chemical engineering of actin filaments within living plant cells.” ChemBioChem 12(1): 132–137.

Elkin, S. R., N. W. Oswald, D. K. Reed, M. Mettlen, J. B. MacMillan and S. L. Schmid (2016). “Ikarugamycin: A Natural Product Inhibitor of Clathrin-Mediated Endocytosis.” Traffic 17(10): 1139–1149.

Guo, B., J. Itami, K. Oikawa, Y. Motoda, T. Kigawa and K. Numata (2019). “Native protein delivery into rice callus using ionic complexes of protein and cell-penetrating peptides.” PLOS ONE 14(7): e0214033.

Hamada, H., Y. Liu, Y. Nagira, R. Miki, N. Taoka and R. Imai (2018). “Biolistic-delivery-based transient CRISPR/Cas9 expression enables in planta genome editing in wheat.” Scientific Reports 8(1): 14422.

Heinicke, E., U. Kumar and D. G. Munoz (1992). “Quantitative dot-blot assay for proteins using enhanced chemiluminescence.” Journal of Immunological Methods 152(2): 227–236.

Hellens, R. P., E. A. Edwards, N. R. Leyland, S. Bean and P. M. Mullineaux (2000). “pGreen: a versatile and flexible binary Ti vector for Agrobacterium-mediated plant transformation.” Plant Molecular Biology 42(6): 819–832.

Hemmer, C., S. Djennane, L. Ackerer, K. Hleibieh, A. Marmonier, S. Gersch, S. Garcia, E. Vigne, V. Komar, M. Perrin, C. Gertz, L. Belval, F. Berthold, B. Monsion, C. Schmitt-Keichinger, O. Lemaire, B. Lorber, C. Gutiérrez, S. Muyldermans, G. Demangeat and C. Ritzenthaler (2018). “Nanobody-mediated resistance to Grapevine fanleaf virus in plants.” Plant Biotechnology Journal 16(2): 660–671.

Hicks, G. R., M. S. S. Harley, M. Shieh and N. V. Raikhel (1995). “Three Classes of Nuclear Import Signald Bind to Plant Nuclei.” Plant Physiology 107(4): 1055–1058.

Hu, C.-D., Y. Chinenov and T. K. Kerppola (2002). “Visualization of Interactions among bZIP and Rel Family Proteins in Living Cells Using Bimolecular Fluorescence Complementation.” Molecular Cell 9(4): 789–798.

Ikeda, M., N. Mitsuda and M. Ohme-Takagi (2009). “Arabidopsis WUSCHEL Is a Bifunctional Transcription Factor That Acts as a Repressor in Stem Cell Regulation and as an Activator in Floral Patterning.” The Plant Cell 21(11): 3493–3505.

Jinek, M., K. Chylinski, I. Fonfara, M. Hauer, J. A. Doudna and E. Charpentier (2012). “A programmable dual-RNA-guided DNA endonuclease in adaptive bacterial immunity.” Science (New York, N.Y.) 337(6096): 816–821.

Kamiyama, D., S. Sekine, B. Barsi-Rhyne, J. Hu, B. Chen, L. A. Gilbert, H. Ishikawa, M. D. Leonetti, W. F. Marshall, J. S. Weissman and B. Huang (2016). “Versatile protein tagging in cells with split fluorescent protein.” Nature Communications 7(1): 11046.

Kosuge, M., T. Takeuchi, I. Nakase, A. T. Jones and S. Futaki (2008). “Cellular Internalization and Distribution of Arginine-Rich Peptides as a Function of Extracellular Peptide Concentration, Serum, and Plasma Membrane Associated Proteoglycans.” Bioconjugate Chemistry 19(3): 656–664.

Kruskal, W. H. and W. A. Wallis (1952). “Use of Ranks in One-Criterion Variance Analysis.” Journal of the American Statistical Association 47(260): 583–621.

Lacroix, A., E. Vengut-Climent, D. de Rochambeau and H. F. Sleiman (2019). “Uptake and Fate of Fluorescently Labeled DNA Nanostructures in Cellular Environments: A Cautionary Tale.” ACS Central Science 5(5): 882–891.

Langel, Ü. (2011). Cell-penetrating peptides, Springer.

Leibfried, A., J. P. C. To, W. Busch, S. Stehling, A. Kehle, M. Demar, J. J. Kieber and J. U. Lohmann (2005). “WUSCHEL controls meristem function by direct regulation of cytokinin-inducible response regulators.” Nature 438(7071): 1172–1175.

Li, H., J. Li, J. Chen, L. Yan and L. Xia (2020). “Precise Modifications of Both Exogenous and Endogenous Genes in Rice by Prime Editing.” Molecular Plant 13(5): 671–674.

Livak, K. J. and T. D. Schmittgen (2001). “Analysis of relative gene expression data using real-time quantitative PCR and the 2(-Delta Delta C(T)) Method.” Methods 25(4): 402–408.

Lowe, K., E. Wu, N. Wang, G. Hoerster, C. Hastings, M.-J. Cho, C. Scelonge, B. Lenderts, M. Chamberlin, J. Cushatt, L. Wang, L. Ryan, T. Khan, J. Chow-Yiu, W. Hua, M. Yu, J. Banh, Z. Bao, K. Brink, E. Igo, B. Rudrappa, P. Shamseer, W. Bruce, L. Newman, B. Shen, P. Zheng, D. Bidney, C. Falco, J. Register, Z.-Y. Zhao, D. Xu, T. Jones and W. Gordon-Kamm (2016). “Morphogenic Regulators Baby boom and Wuschel Improve Monocot Transformation.” The Plant Cell 28(9): 1998–2015.

Martin-Ortigosa, S., D. J. Peterson, J. S. Valenstein, V. S.-Y. Lin, B. G. Trewyn, L. A. Lyznik and K. Wang (2013). “Mesoporous Silica Nanoparticle-Mediated Intracellular Cre Protein Delivery for Maize Genome Editing via loxP Site Excision,” Plant Physiology 164(2): 537–547.

Martin, K., K. Kopperud, R. Chakrabarty, R. Banerjee, R. Brooks and M. M. Goodin (2009). “Transient expression in Nicotiana benthamiana fluorescent marker lines provides enhanced definition of protein localization, movement and interactions in planta.” The Plant Journal 59(1): 150–162.

Martin, R. M., G. Ter-Avetisyan, H. D. Herce, A. K. Ludwig, G. Lättig-Tünnemann and M. C. Cardoso (2015). “Principles of protein targeting to the nucleolus.” Nucleus 6(4): 314–325.

Milech, N., B. A. Longville, P. T. Cunningham, M. N. Scobie, H. M. Bogdawa, S. Winslow, M. Anastasas, T. Connor, F. Ong and S. R. Stone (2015). “GFP-complementation assay to detect functional CPP and protein delivery into living cells.” Scientific reports 5(1): 1–11.

Numata, K., Y. Horii, K. Oikawa, Y. Miyagi, T. Demura and M. Ohtani (2018). “Library screening of cell-penetrating peptide for BY-2 cells, leaves of Arabidopsis, tobacco, tomato, poplar, and rice callus.” Scientific Reports 8(1): 10966.

Otsu, N. (1979). “A Threshold Selection Method from Gray-Level Histograms.” IEEE Transactions on Systems, Man, and Cybernetics 9(1): 62–66.

Palm, C., M. Jayamanne, M. Kjellander and M. Hällbrink (2007). “Peptide degradation is a critical determinant for cell-penetrating peptide uptake.” Biochimica et Biophysica Acta (BBA)-Biomembranes 1768(7): 1769–1776.

Pantarotto, D., J. P. Briand, M. Prato and A. Bianco (2004). “Translocation of bioactive peptides across cell membranes by carbon nanotubes.” Chem Commun (Camb)(1): 16–17.

Park, E., H.-Y. Lee, J. Woo, D. Choi and S. P. Dinesh-Kumar (2017). “Spatiotemporal Monitoring of <EM>Pseudomonas syringae</EM> Effectors via Type III Secretion Using Split Fluorescent Protein Fragments.” The Plant Cell 29(7): 1571–1584.

Patel, S. G., E. J. Sayers, L. He, R. Narayan, T. L. Williams, E. M. Mills, R. K. Allemann, L. Y. P. Luk, A. T. Jones and Y.-H. Tsai (2019). “Cell-penetrating peptide sequence and modification dependent uptake and subcellular distribution of green florescent protein in different cell lines.” Scientific Reports 9(1): 6298.

Pawley, J. B. (2006). Handbook of Biological Confocal Microscopy. Boston, MA, Springer.

Peraro, L., K. L. Deprey, M. K. Moser, Z. Zou, H. L. Ball, B. Levine and J. A. Kritzer (2018). “Cell Penetration Profiling Using the Chloroalkane Penetration Assay.” Journal of the American Chemical Society 140(36): 11360–11369.

Riedl, J., A. H. Crevenna, K. Kessenbrock, J. H. Yu, D. Neukirchen, M. Bista, F. Bradke, D. Jenne, T. A. Holak, Z. Werb, M. Sixt and R. Wedlich-Soldner (2008). “Lifeact: a versatile marker to visualize F-actin.” Nature methods 5(7): 605–607.

Sarrion-Perdigones, A., M. Vazquez-Vilar, J. Palací, B. Castelijns, J. Forment, P. Ziarsolo, J. Blanca, A. Granell and D. Orzaez (2013). “GoldenBraid 2.0: A Comprehensive DNA Assembly Framework for Plant Synthetic Biology.” Plant Physiology 162(3): 1618–1631.

Schmidt, S., M. J. W. Adjobo-Hermans, R. Wallbrecher, W. P. R. Verdurmen, P. H. M. Bovée-Geurts, J. van Oostrum, F. Milletti, T. Enderle and R. Brock (2015). “Detecting Cytosolic Peptide Delivery with the GFP Complementation Assay in the Low Micromolar Range.” Angewandte Chemie International Edition 54(50): 15105–15108.

Schoof, H., M. Lenhard, A. Haecker, K. F. X. Mayer, G. Jürgens and T. Laux (2000). “The Stem Cell Population of Arabidopsis Shoot Meristems Is Maintained by a Regulatory Loop between the CLAVATA and WUSCHEL Genes.” Cell 100(6): 635–644.

Schwartz, S. H., B. Hendrix, P. Hoffer, R. A. Sanders and W. Zheng (2020). “Carbon Dots for Efficient Small Interfering RNA Delivery and Gene Silencing in Plants.” Plant Physiology 184(2): 647–657.

Serna, L. (2005). “A simple method for discriminating between cell membrane and cytosolic proteins.” New Phytologist 165(3): 947–952.

Stirling, D. R., M. J. Swain-Bowden, A. M. Lucas, A. E. Carpenter, B. A. Cimini and A. Goodman (2021). “CellProfiler 4: improvements in speed, utility and usability.” BMC Bioinformatics 22(1): 433.

Sugiura, D., I. Terashima and J. R. Evans (2020). “A Decrease in Mesophyll Conductance by Cell-Wall Thickening Contributes to Photosynthetic Downregulation.” Plant Physiology 183(4): 1600–1611.

Teo, S. L. Y., J. J. Rennick, D. Yuen, H. Al-Wassiti, A. P. R. Johnston and C. W. Pouton (2021). “Unravelling cytosolic delivery of cell penetrating peptides with a quantitative endosomal escape assay.” Nature Communications 12(1): 3721.

Thagun, C., Y. Horii, M. Mori, S. Fujita, M. Ohtani, K. Tsuchiya, Y. Kodama, M. Odahara and K. Numata (2022). “Non-transgenic Gene Modulation via Spray Delivery of Nucleic Acid/Peptide Complexes into Plant Nuclei and Chloroplasts.” ACS Nano.

Wallbrecher, R., T. Ackels, R. A. Olea, M. J. Klein, L. Caillon, J. Schiller, P. H. Bovée-Geurts, T. H. van Kuppevelt, A. S. Ulrich, M. Spehr, M. J. W. Adjobo-Hermans and R. Brock (2017). “Membrane permeation of arginine-rich cell-penetrating peptides independent of transmembrane potential as a function of lipid composition and membrane fluidity.” Journal of Controlled Release 256: 68–78.

Wallmann, A. and C. Kesten (2020). “Common Functions of Disordered Proteins across Evolutionary Distant Organisms.” International Journal of Molecular Sciences 21(6): 2105.

Wang, J. W., F. J. Cunningham, N. S. Goh, N. N. Boozarpour, M. Pham and M. P. Landry (2021). “Nanoparticles for protein delivery in planta.” Current Opinion in Plant Biology 60: 102052.

Wang, R. and M. G. Brattain (2007). “The maximal size of protein to diffuse through the nuclear pore is larger than 60 kDa.” FEBS Letters 581(17): 3164–3170.

Wang, Y., Y. Wang and Y. Wang (2020). “Apoplastic Proteases: Powerful Weapons against Pathogen Infection in Plants.” Plant Communications 1(4): 100085.

Wright, K. M., N. T. Wood, A. G. Roberts, S. Chapman, P. Boevink, K. M. MacKenzie and K. J. Oparka (2007). “Targeting of TMV Movement Protein to Plasmodesmata Requires the Actin/ER Network; Evidence From FRAP.” Traffic 8(1): 21–31.

Xu, X., A. Chemparathy, L. Zeng, H. R. Kempton, S. Shang, M. Nakamura and L. S. Qi (2021). “Engineered miniature CRISPR-Cas system for mammalian genome regulation and editing.” Molecular Cell 81(20): 4333–4345.e4334.

Youngblood, D. S., S. A. Hatlevig, J. N. Hassinger, P. L. Iversen and H. M. Moulton (2007). “Stability of Cell-Penetrating Peptide−Morpholino Oligomer Conjugates in Human Serum and in Cells.” Bioconjugate Chemistry 18(1): 50–60.

Yuan, J. S., A. Reed, F. Chen and C. N. Stewart (2006). “Statistical analysis of real-time PCR data.” BMC Bioinformatics 7(1): 85.

Zhang, H., G. S. Demirer, H. Zhang, T. Ye, N. S. Goh, A. J. Aditham, F. J. Cunningham, C. Fan and M. P. Landry (2019). “DNA nanostructures coordinate gene silencing in mature plants.” Proceedings of the National Academy of Sciences 116(15): 7543.

Zhang, H., N. S. Goh, J. W. Wang, R. L. Pinals, E. González-Grandío, G. S. Demirer, S. Butrus, S. C. Fakra, A. Del Rio Flores, R. Zhai, B. Zhao, S.-J. Park and M. P. Landry (2022). “Nanoparticle cellular internalization is not required for RNA delivery to mature plant leaves.” Nature Nanotechnology 17(2): 197–205.

Ziegler, A., P. Nervi, M. Dürrenberger and J. Seelig (2005). “The Cationic Cell-Penetrating Peptide CPPTAT Derived from the HIV-1 Protein TAT Is Rapidly Transported into Living Fibroblasts:!] Optical, Biophysical, and Metabolic Evidence.” Biochemistry 44(1): 138–148.

Zuo, J., Q.-W. Niu, G. Frugis and N.-H. Chua (2002). “The WUSCHEL gene promotes vegetative-to-embryonic transition in Arabidopsis.” The Plant Journal 30(3): 349–359.

