## Supplemental Figures for "Fluorescence complementation enables quantitative imaging of cell penetrating peptide-mediated protein delivery in plants including WUSCHEL transcription factor"

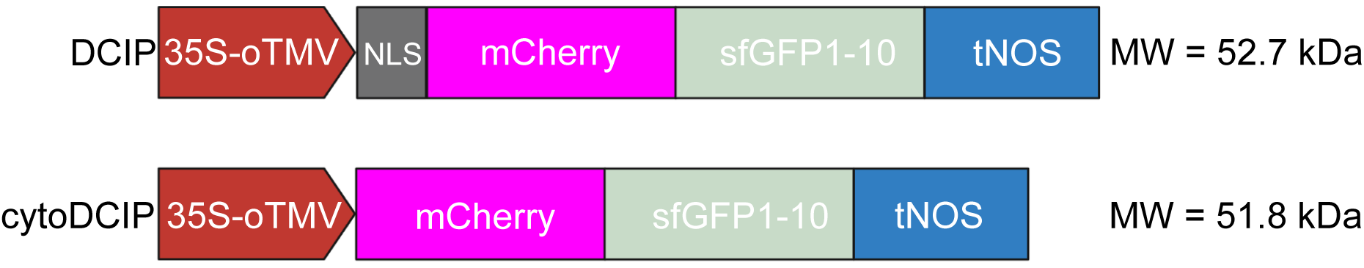

**Supplemental Figure 1.** Schematic of DCIP and cytoDCIP transcriptional unit. Expression is driven by a 35s promoter and terminated by tNOS. DCIP possess a SV40 NLS for nuclear localization whereas cytoDCIP does not and localizes to the cytosol. Both vectors were constructed as level-1 assemblies in Goldenbraid 2.0.

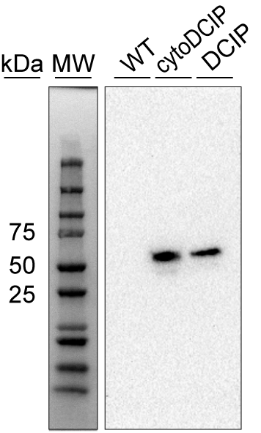

**Supplemental Figure 2.** Western Blot using anti-mCherry primary antibody and *N. benthamiana* leaf lysates 3 d.p.i. with either DCIP or cytoDCIP showing both protein fusions at the predicted molecular weight.

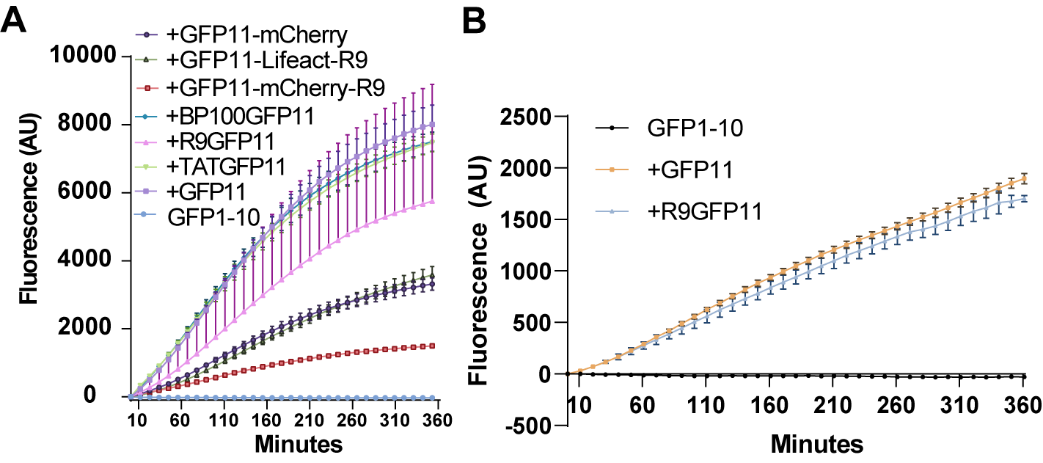

**Supplemental Figure 3.** *In vitro* GFP complementation assay using recombinantly expressed and purified sfGFP1-10 and all GFP11 containing constructs used in this study. A final concentration of 5μM GFP1-10 his-tag eluate and 10μM of GFP11 containing protein or peptide were incubated for up to 6 hours. Fluorescence was measured every minute on a Biorad CFX96 qPCR machine set at (A) 21°C or (B) 4°C.

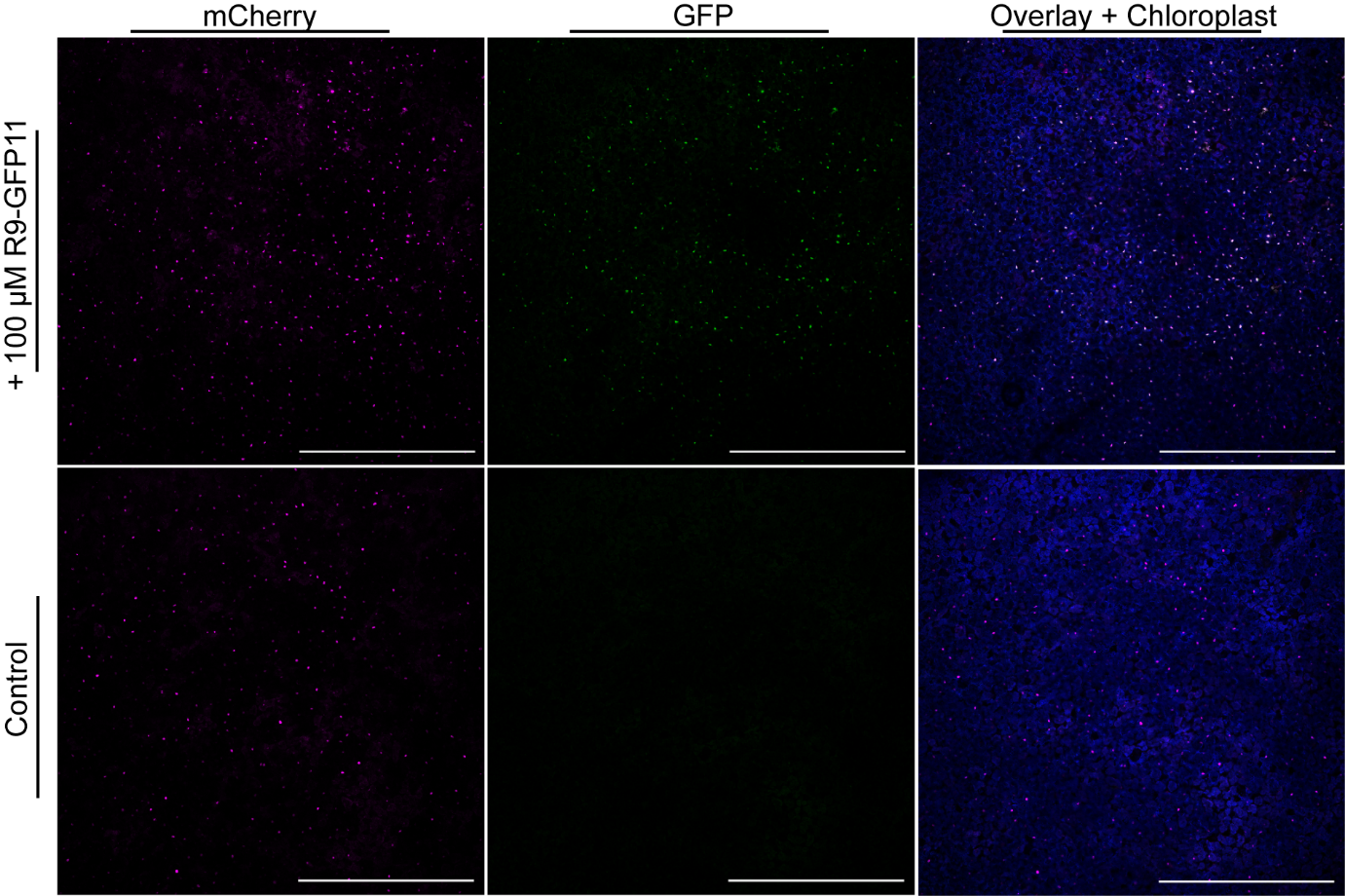

**Supplemental Figure 4.** DCIP expressing plant infiltrated with either 100μM R9-GFP11 (above) or water control (below) and imaged using confocal microscopy at 4-5H using a 5x objective. mCherry fluorescence is pseudocolored magenta and sfGFP fluorescence is colored green. Chloroplast autofluorescence is colored blue. Scale bar is 1mm.

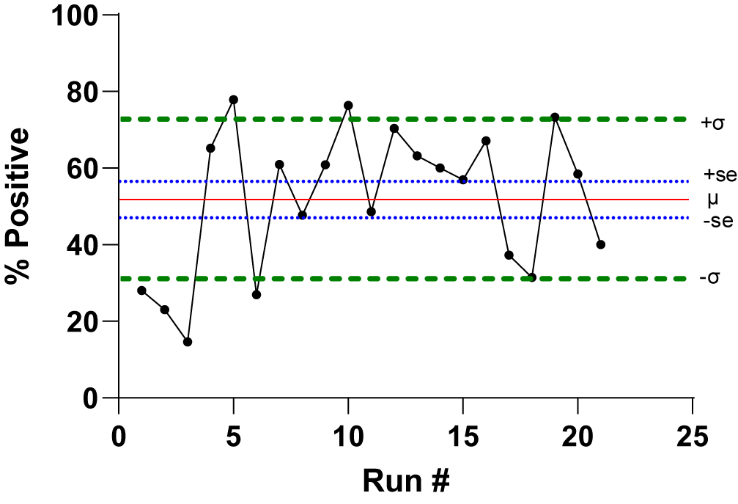

**Supplemental Figure 5.** Percentage of GFP positive nuclei in DCIP expressing *N. benthamiana* treated with 100μM R9-GFP11 for 4-5H in the first 21 DCIP experiments performed. Data were pooled across several experiments to probe the innate variability in R9 delivery across multiple plants. Standard error and standard deviation are marked with a dotted and dashed line respectively. The mean, 52%, is marked with a solid line.

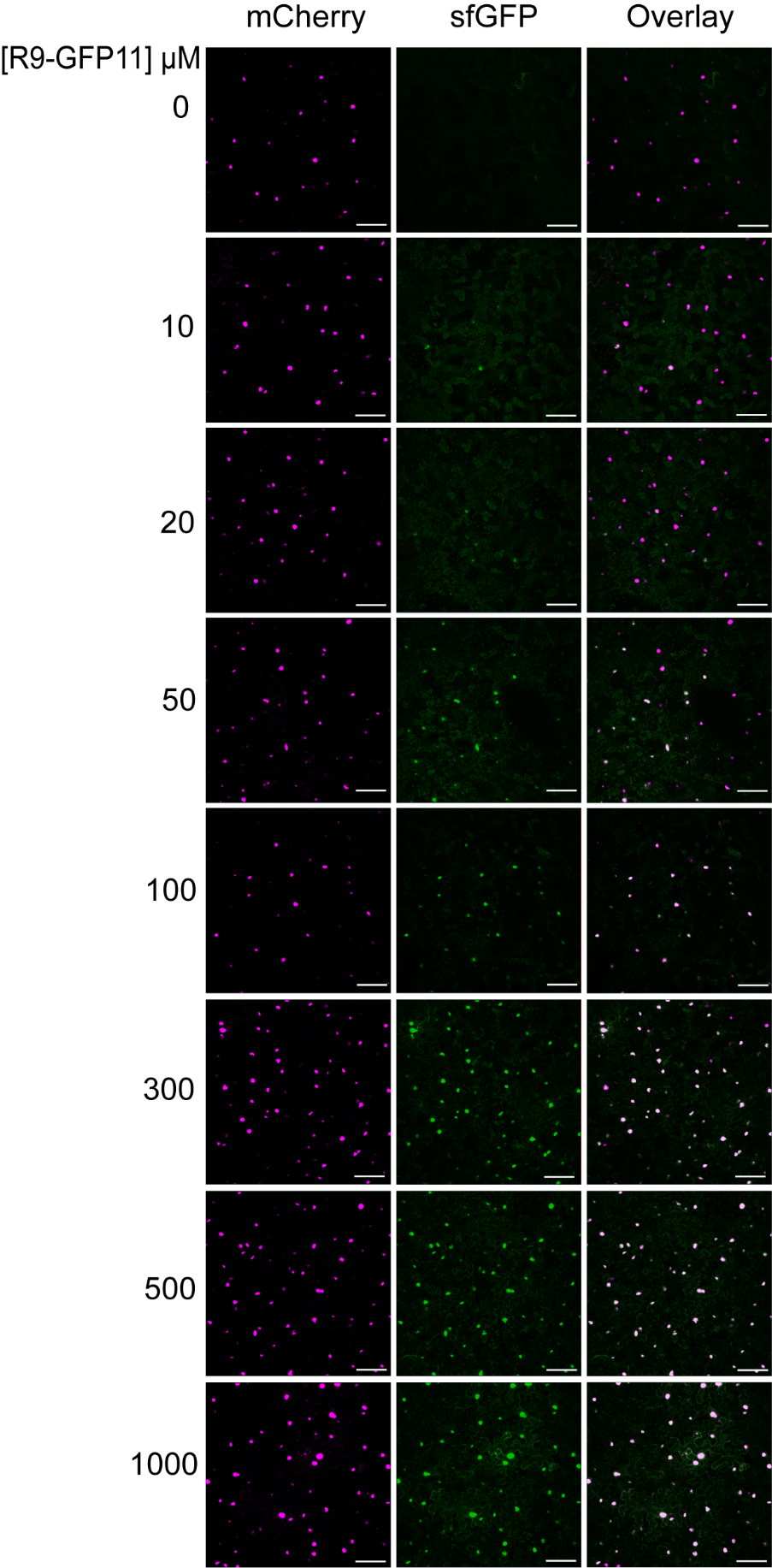
 **Supplemental Figure 6.** Representative two-color maximum intensity projections of DCIP expressing leaves infiltrated with 0-1000 µM R9-GFP11 and incubated for 4-5 hours. Scale bar is 100μm. mCherry is pseudocolored magenta and sfGFP is pseudocolored green. Overlay results in white coloration

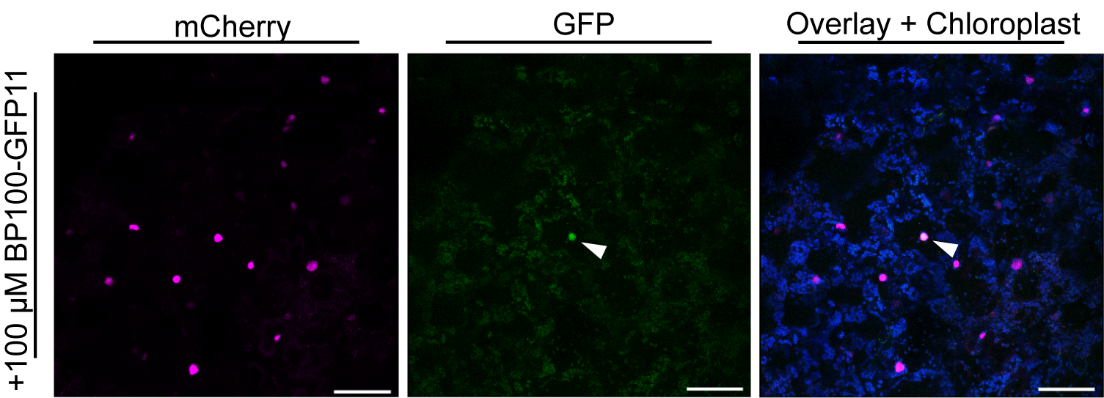

**Supplemental Figure 7.** Example maximum intensity projection of sfGFP complementation as resulting from 100μM BP100-GFP11 incubation in a DCIP expressing leaf disc for 4-5H. Scale bar is 100μm. sfGFP fluorescent nucleus is marked by a white triangle. mCherry is pseudocolored magenta, sfGFP is pseudocolored green, and chloroplast autofluorescence is pseudocolored blue. Overlay results in white coloration.

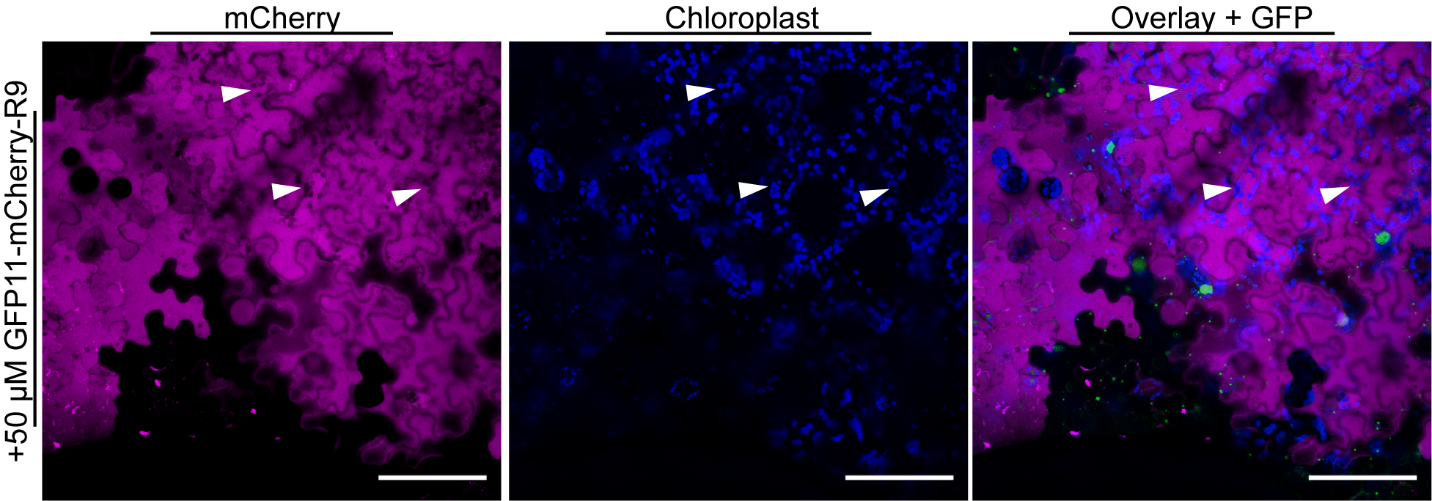

**Supplemental Figure 8.** Single confocal imaging slice at 5H after infiltration with 50 μM GFP11-mCherry-R9 into a DCIP expressing *N. benthamiana* leaf. White triangles mark excluded regions in the mCherry fluorescence channel (magenta) and the chloroplast autofluorescence (blue) channels. Both channels overlaid with sfGFP fluorescence (green, right image). Scale bar is 100 μM.

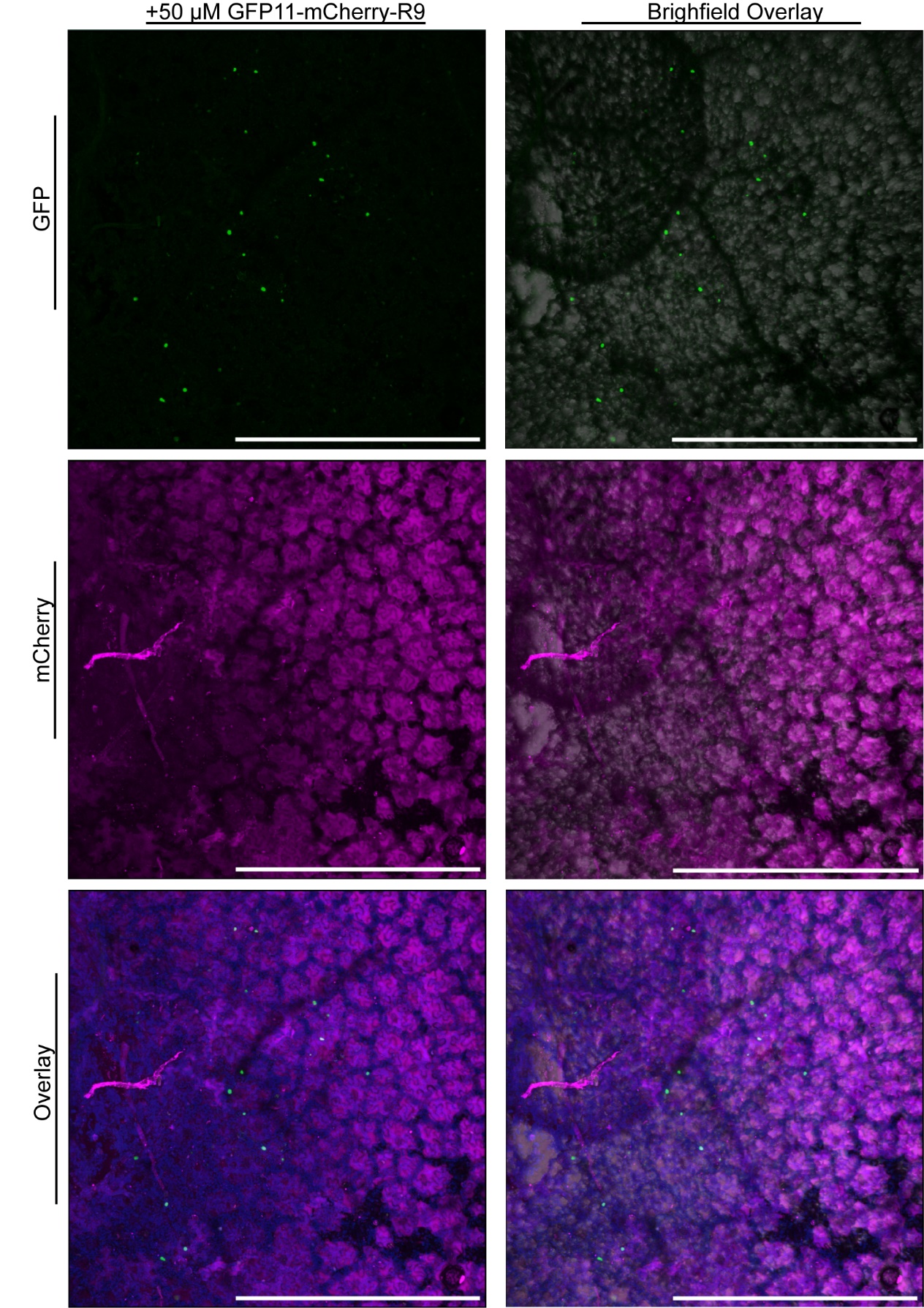

**Supplemental Figure 9.** DCIP expressing plant infiltrated with 50μM GFP11-mCherry-R9 and imaged using confocal microscopy at 5H using a 5x objective. mCherry fluorescence is pseudocolored magenta and sfGFP fluorescence is colored green. Green nuclear fluorescence resulting from successful delivery appear as small circular objects. Chloroplast autofluorescence is colored blue. Transmitted light is overlayed in the right column to show anatomy. Scale bar is 1mm.

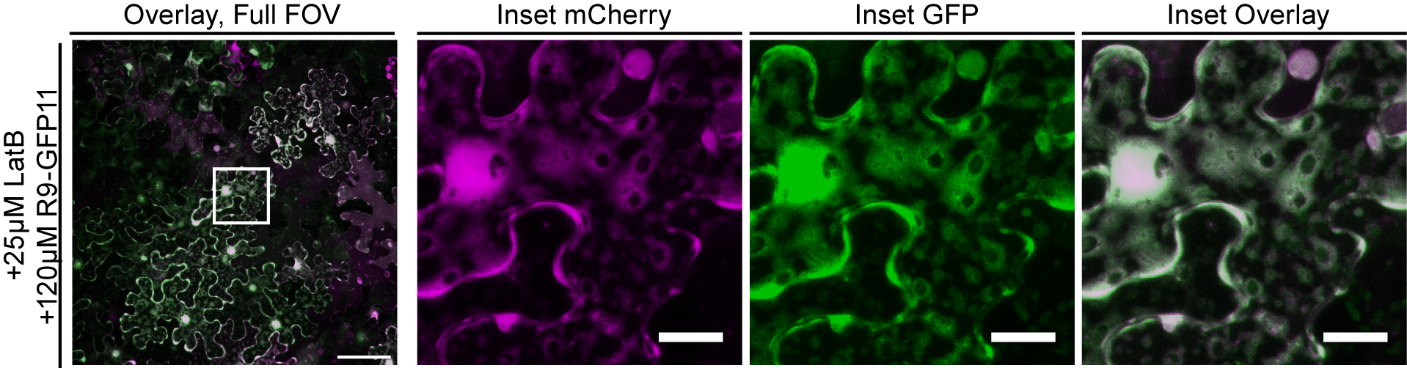

**Supplemental Figure 10.** Representative confocal micrograph (Standard Deviation Projection) of a leaf disc from a cytoDCIP expressing leaf infiltrated with 120 µM R9-GFP11 and 25µM LatB after 6H of incubation. Full FOV scale bar is 100 µm; inset is 20 µm. mCherry is pseudocolored magenta and sfGFP pseudocolored green.

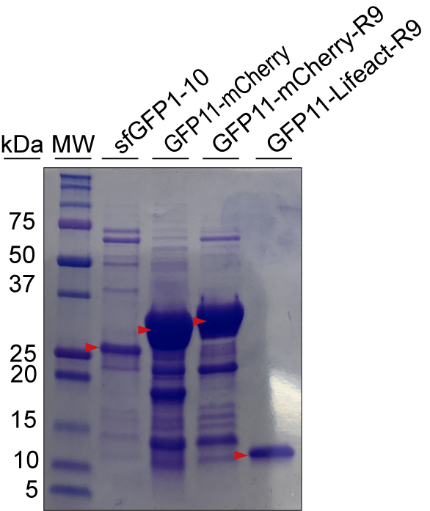

**Supplemental Figure 11.** SDS-PAGE of first four recombinant proteins used in this study. Red triangles indicate the protein of interest. Proteins were stained with Coomassie R-250.

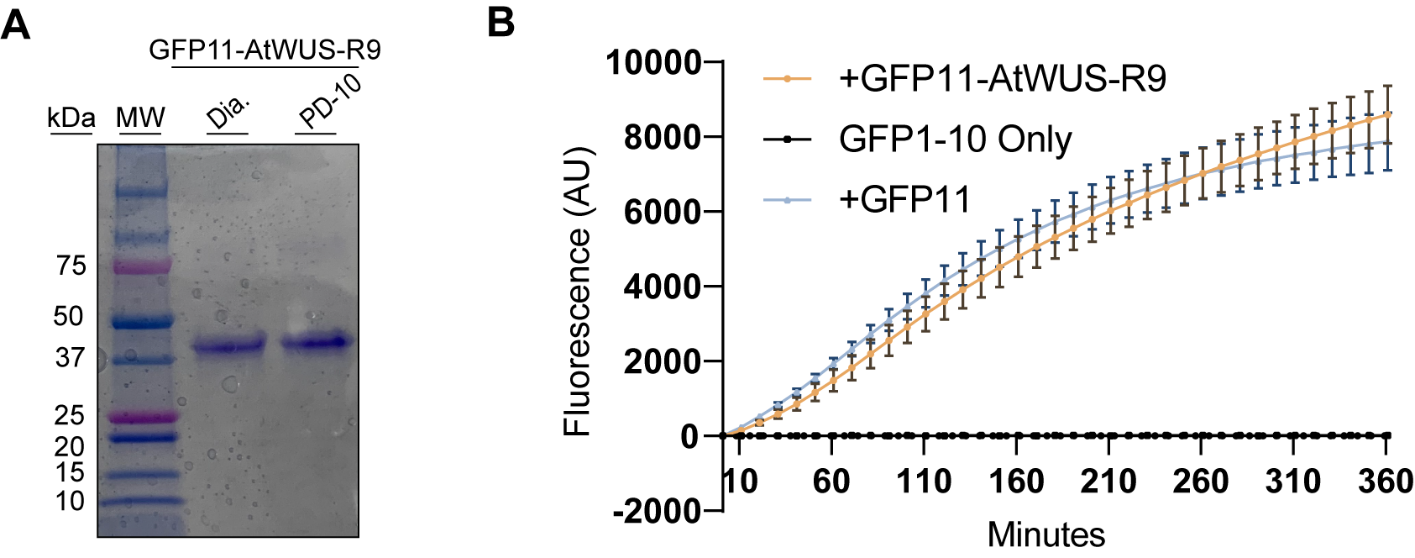

**Supplemental Figure 12.** A, SDS-PAGE of purified GFP11-AtWUS-R9 exchanged into buffer P by dialysis or by desalting column. B, *In vitro* complementation assay of GFP11-AtWUS-R9. A final concentration of 6µM test protein and 4µM GFP1-10 in addition to 200mM Tris, pH 8.0 was used for this assay.
