## Supplemental Tables for "Fluorescence complementation enables quantitative imaging of cell penetrating peptide-mediated protein delivery in plants including WUSCHEL transcription factor"

**Supplemental Table S1.** Plasmids produced or utilized for cloning.

| **Plasmid Name** | **Notes** | **Source** | **Protein Product Sequence** | **Protein Product Length (AA)** | **Protein Product MW (kDa)** |
| --- | --- | --- | --- | --- | --- |
| DCIP | 35s::NLS:mCherry:sfGFP1-10::tNOS fusion for nuclear localized expression using agrobacterium in pDGB1-a1R vector backbone | Generated in this work | MPKKKRKVGVSKGEEDNMAIIKEFMRFKVHMEGSVNGHEFEIEGEGEGRPYEGTQTAKLKVTKGGPLPFAWDILSPQFMYGSKAYVKHPADIPDYLKLSFPEGFKWERVMNFEDGGVVTVTQDSSLQDGEFIYKVKLRGTNFPSDGPVMQKKTMGWEASSERMYPEDGALKGEIKQRLKLKDGGHYDAEVKTTYKAKKPVQLPGAYNVNIKLDITSHNEDYTIVEQYERAEGRHSTGGMDELYKGGGGSGGGSLQMIDSKGEELFTGVVPILVELDGDVNGHKFSVRGEGEGDATIGKLTLKFICTTGKLPVPWPTLVTTLTYGVQCFSRYPDHMKRHDFFKSAMPEGYVQERTISFKDDGKYKTRAVVKFEGDTLVNRIELKGTDFKEDGNILGHKLEYNFNSHNVYITADKQKNGIKANFTVRHNVEDGSVQLADHYQQNTPIGDGPVLLPDNHYLSTQTVLSKDPNEK* | 471 | 52.7 |
| cytoCDCIP | 35s::mCherry:sfGFP1-10::tNOS fusion for cytosolic expression using agrobacterium in pDGB1-a1R vector backbone | Generated in this work | MVSKGEEDNMAIIKEFMRFKVHMEGSVNGHEFEIEGEGEGRPYEGTQTAKLKVTKGGPLPFAWDILSPQFMYGSKAYVKHPADIPDYLKLSFPEGFKWERVMNFEDGGVVTVTQDSSLQDGEFIYKVKLRGTNFPSDGPVMQKKTMGWEASSERMYPEDGALKGEIKQRLKLKDGGHYDAEVKTTYKAKKPVQLPGAYNVNIKLDITSHNEDYTIVEQYERAEGRHSTGGMDELYKGGGGSGGGSLQMIDSKGEELFTGVVPILVELDGDVNGHKFSVRGEGEGDATIGKLTLKFICTTGKLPVPWPTLVTTLTYGVQCFSRYPDHMKRHDFFKSAMPEGYVQERTISFKDDGKYKTRAVVKFEGDTLVNRIELKGTDFKEDGNILGHKLEYNFNSHNVYITADKQKNGIKANFTVRHNVEDGSVQLADHYQQNTPIGDGPVLLPDNHYLSTQTVLSKDPNEK* | 463 | 51.8 |
| 1B-GFP1-10 | sfGFP1-10 inserted into 1B for E. coli expression and Hisx6 tag | Generated in this work | MGSSHHHHHHENLYFQSNAMIDSKGEELFTGVVPILVELDGDVNGHKFSVRGEGEGDATIGKLTLKFICTTGKLPVPWPTLVTTLTYGVQCFSRYPDHMKRHDFFKSAMPEGYVQERTISFKDDGKYKTRAVVKFEGDTLVNRIELKGTDFKEDGNILGHKLEYNFNSHNVYITADKQKNGIKANFTVRHNVEDGSVQLADHYQQNTPIGDGPVLLPDNHYLSTQTVLSKDPNEKGIGSG* | 240 | 26.9 |
| 1BR9-mCherry | mCherry insertion of 1BR9 for E. coli expression and Hisx6 tag | Generated in this work | MGSSHHHHHHENLYFQSNARDHMVLHEYVNAAGITGGGGSGGGGSYFQSNAVSKGEEDNMAIIKEFMRFKVHMEGSVNGHEFEIEGEGEGRPYEGTQTAKLKVTKGGPLPFAWDILSPQFMYGSKAYVKHPADIPDYLKLSFPEGFKWERVMNFEDGGVVTVTQDSSLQDGEFIYKVKLRGTNFPSDGPVMQKKTMGWEASSERMYPEDGALKGEIKQRLKLKDGGHYDAEVKTTYKAKKPVQLPGAYNVNIKLDITSHNEDYTIVEQYERAEGRHSTGGMDELYKGIGSGSNGSSGSVSRRRRRRRRR* | 309 | 34.5 |
| 1BR9-mCherrySTOP | mCherry insertion of 1BR9 E. coli expression and Hisx6 tag with stop codon to prevent R9 tagging | Generated in this work | MGSSHHHHHHENLYFQSNARDHMVLHEYVNAAGITGGGGSGGGGSYFQSNAVSKGEEDNMAIIKEFMRFKVHMEGSVNGHEFEIEGEGEGRPYEGTQTAKLKVTKGGPLPFAWDILSPQFMYGSKAYVKHPADIPDYLKLSFPEGFKWERVMNFEDGGVVTVTQDSSLQDGEFIYKVKLRGTNFPSDGPVMQKKTMGWEASSERMYPEDGALKGEIKQRLKLKDGGHYDAEVKTTYKAKKPVQLPGAYNVNIKLDITSHNEDYTIVEQYERAEGRHSTGGMDELYK | 286 | 32.0 |
| 1BR9-Lifeact | Lifeact peptide insertion of 1BR9 E. coli expression and Hisx6 tag | Generated in this work | MGSSHHHHHHENLYFQSNARDHMVLHEYVNAAGITGGGGSGGGGSYFQSNAMGVADLIKKFESISKEEGIGSGSNGSSGSVSRRRRRRRRR* | 91 | 9.87 |
| PEP101E-NLS-mCherry | UBQ10 driven expression of NLS-mCherry-sfGFP1-10 and Hisx6 tag | Generated in this work | MPKKKRKVGVSKGEEDNMAIIKEFMRFKVHMEGSVNGHEFEIEGEGEGRPYEGTQTAKLKVTKGGPLPFAWDILSPQFMYGSKAYVKHPADIPDYLKLSFPEGFKWERVMNFEDGGVVTVTQDSSLQDGEFIYKVKLRGTNFPSDGPVMQKKTMGWEASSERMYPEDGALKGEIKQRLKLKDGGHYDAEVKTTYKAKKPVQLPGAYNVNIKLDITSHNEDYTIVEQYERAEGRHSTGGMDELYKGGGGSGGGSLQMIDSKGEELFTGVVPILVELDGDVNGHKFSVRGEGEGDATIGKLTLKFICTTGKLPVPWPTLVTTLTYGVQCFSRYPDHMKRHDFFKSAMPEGYVQERTISFKDDGKYKTRAVVKFEGDTLVNRIELKGTDFKEDGNILGHKLEYNFNSHNVYITADKQKNGIKANFTVRHNVEDGSVQLADHYQQNTPIGDGPVLLPDNHYLSTQTVLSKDPNEK* | 471 | 52.7 |
| 1BR9 | LIC backbone derivatized from 1B for expression of N-terminal GFP11 and C-terminal R9 tagged proteins in E. coli | Generated in this work | MGSSHHHHHHENLYFQSNARDHMVLHEYVNAAGITGGGGSGGGGSYFQSN(A(yORF)G)IGSGSNGSSGSVSRRRRRRRRR* | 72 | 7.83 |
| PEP101E | Cytosolic UBQ10 driven expression of sfGFP1-10 Plasmid from Prof. Dinesh Kumar (Park et al Plant Cell. 2017 Jul;29(7):1571-1584.) | Addgene #97387 |  |  |  |
| 1B | LIC backbone for hisx6 tagged expression of proteins in E. coli from Scott Gradia at UC Berkeley MacroLab | Addgene #29653 |  |  |  |
| H6-mCherry | mCherry coding sequeince from Scott Gradia at UC Berkeley MacroLab | Addgene #29722 |  |  |  |
| 1BR9-AtWUS | E. coli codon optimized AtWUS insertion of 1BR9 for E. coli expression and Hisx6 tag | Generated in this work | MGSSHHHHHHENLYFQSNARDHMVLHEYVNAAGITGGGGSGGGGSYFQSNAEPPQHQHHHHQADQESGNNNNNKSGSGGYTCRQTSTRWTPTTEQIKILKELYYNNAIRSPTADQIQKITARLRQFGKIEGKNVFYWFQNHKARERQKKRFNGTNMTTPSSSPNSVMMAANDHYHPLLHHHHGVPMQRPANSVNVKLNQDHHLYHHNKPYPSFNNGNLNHASSGTECGVVNASNGYMSSHVYGSMEQDCSMNYNNVGGGWANMDHHYSSAPYNFFDRAKPLFGLEGHQEEEECGGDAYLEHRRTLPLFPMHGEDHINGGSGAIWKYGQSEVRPCASLELRLNGIGSGSNGSSGSVSRRRRRRRRR | 365 | 41 |

**Supplemental Table S2** DNA sequences and primers used.

| **DNA Set** | **FW 5->3' Sequence** | **REV 5->3' Sequence** | **Use** |
| --- | --- | --- | --- |
| 1 | AAACTGCAGATGCCAAAGAAAAAGCGGAAAGTCGGAGTGAGCAAGGGCGAGGAG | AAACTGCAGGCTGCCACCGCCGCTACCGCCACCGCCCTTGTACAGCTCGTCCATGC | Amplification of mCherry from h6 mCherry Vector with PSTI overhangs for 5' insertion into Dinesh-kumar sfGFP1-10 vector with an NLS sequence and disordered linker |
| 2 | TACTTCCAATCCAATGCAGTGAGCAAGGGCGAGGAG | CTCCCACTACCAATGCCTTACTTGTACAGCTCGTCCATG | Amplification of mCherry from h6 mCherry Vector for 1BR9 cloning (with stop codon) |
| 3 | TACTTCCAATCCAATGCAGTGAGCAAGGGCGAGGAG | CTCCCACTACCAATGCCCTTGTACAGCTCGTCCATG | Amplification of mCherry from h6 mCherry Vector for 1BR9 cloning (no stop codon) |
| 4 | TACTTCCAATCCAATGCCACCATGATCGATAGCAAAGGAGAAG | TTATCCACTTCCAATGTTATTATCACTTTTCGTTGGGATCTTTC | Amplification of sfGFP1-10 from plasmid from Dinesh-kumar for cloning into 1B |
| 5 | GCGCCGTCTCGCTCAAAGCTCACTTTT | GCGCCGTCTCGCTCGAATGCCAAAGA | Amplification of NLS:mCherry:GFP1-10 for goldenbraid cloning into pUPD2 |
| 6 | GCGCCGTCTCGCTCGAATGGTGAGCA | GCGCCGTCTCGCTCGAATGCCAAAGA | Amplification of mCherry:GFP1-10 for goldenbraid cloning into pUPD2 |
| 7 | TACTTCCAATCCAATGCACGCGATCACATGGTCCTGCACGAGTACGTGAACGCCGCCGGGATCACTGGTGGCGGAGGTTCTGGAGGCGGTGGATCGTACTTCCAATCCAATATTGGTAGTGGGAGCAACGGCAGCAGCGGATCCGTGAGCCGCCGTCGCCGTCGCCGTCGCCGTCGCTAATAACATTGGAAGTGGATAA | TTATCCACTTCCAATGTTATTAGCGACGGCGACGGCGACGGCGACGGCGGCTCACGGATCCGCTGCTGCCGTTGCTCCCACTACCAATATTGGATTGGAAGTACGATCCACCGCCTCCAGAACCTCCGCCACCAGTGATCCCGGCGGCGTTCACGTACTCGTGCAGGACCATGTGATCGCGTGCATTGGATTGGAAGTA | LIC Insert for conversion of 1B to 1BR9 |
| 8 | TACTTCCAATCCAATGCAATGGGCGTGGCCGACCTGATCAAGAAGTTCGAGAGCATCAGCAAGGAAGAGGGCATTGGTAGTGGGAG | CTCCCACTACCAATGCCCTCTTCCTTGCTGATGCTCTCGAACTTCTTGATCAGGTCGGCCACGCCCATTGCATTGGATTGGAAGTA | Lifeact Insert for LIC into 1BR9 |
| 9 | ATGGAACCTCCCCAGCACCAGCACCATCACCACCAGGCTGATCAAGAGTCGGGAAATAACAACAACAATAAAAGCGGGAGCGGGGGTTACACCTGTCGTCAAACATCCACGCGCTGGACACCAACCACTGAACAAATCAAAATTTTGAAAGAATTATACTATAACAATGCCATTCGTTCACCTACCGCAGATCAAATCCAAAAAATTACAGCACGCTTGCGCCAGTTCGGAAAGATTGAGGGAAAAAACGTGTTCTACTGGTTTCAGAATCATAAGGCACGTGAACGTCAAAAAAAACGTTTTAACGGGACTAATATGACAACACCGAGTTCGAGTCCCAACTCCGTTATGATGGCAGCGAACGATCACTATCATCCATTACTGCACCACCATCATGGAGTCCCAATGCAGCGTCCAGCCAATTCCGTGAATGTCAAGCTGAACCAAGACCACCATCTTTATCACCACAATAAGCCTTACCCATCTTTTAATAACGGCAACCTGAATCACGCCAGCAGTGGCACCGAATGCGGGGTCGTCAATGCGTCGAATGGATATATGTCGTCGCACGTCTACGGTTCTATGGAACAAGACTGTTCAATGAATTACAACAACGTAGGAGGGGGCTGGGCCAATATGGACCATCACTACAGCTCTGCGCCGTACAATTTCTTCGACCGTGCTAAACCGTTATTCGGCTTAGAGGGACATCAGGAGGAGGAGGAATGCGGGGGTGACGCTTACTTAGAACATCGTCGTACTTTACCGCTTTTCCCGATGCATGGAGAGGACCATATCAATGGTGGGTCTGGCGCCATTTGGAAGTACGGACAGTCGGAGGTACGCCCTTGCGCAAGTCTTGAGCTGCGTTTGAATTAG | CTAATTCAAACGCAGCTCAAGACTTGCGCAAGGGCGTACCTCCGACTGTCCGTACTTCCAAATGGCGCCAGACCCACCATTGATATGGTCCTCTCCATGCATCGGGAAAAGCGGTAAAGTACGACGATGTTCTAAGTAAGCGTCACCCCCGCATTCCTCCTCCTCCTGATGTCCCTCTAAGCCGAATAACGGTTTAGCACGGTCGAAGAAATTGTACGGCGCAGAGCTGTAGTGATGGTCCATATTGGCCCAGCCCCCTCCTACGTTGTTGTAATTCATTGAACAGTCTTGTTCCATAGAACCGTAGACGTGCGACGACATATATCCATTCGACGCATTGACGACCCCGCATTCGGTGCCACTGCTGGCGTGATTCAGGTTGCCGTTATTAAAAGATGGGTAAGGCTTATTGTGGTGATAAAGATGGTGGTCTTGGTTCAGCTTGACATTCACGGAATTGGCTGGACGCTGCATTGGGACTCCATGATGGTGGTGCAGTAATGGATGATAGTGATCGTTCGCTGCCATCATAACGGAGTTGGGACTCGAACTCGGTGTTGTCATATTAGTCCCGTTAAAACGTTTTTTTTGACGTTCACGTGCCTTATGATTCTGAAACCAGTAGAACACGTTTTTTCCCTCAATCTTTCCGAACTGGCGCAAGCGTGCTGTAATTTTTTGGATTTGATCTGCGGTAGGTGAACGAATGGCATTGTTATAGTATAATTCTTTCAAAATTTTGATTTGTTCAGTGGTTGGTGTCCAGCGCGTGGATGTTTGACGACAGGTGTAACCCCCGCTCCCGCTTTTATTGTTGTTGTTATTTCCCGACTCTTGATCAGCCTGGTGGTGATGGTGCTGGTGCTGGGGAGGTTCCAT | E. Coli codon optimized DNA sequence for AtWUS |
| 10 | TACTTCCAATCCAATGCAGAACCTCCCCAGCACCAG | CTCCCACTACCAATGCCATTCAAACGCAGCTCAAGAC | Amplification of AtWUS sequence from synethic DNA for insertion into 1BR9 |
| 11 | AACTCTATGCAGCATTTGATCCAC | TGATTGCATATCTTTATCGCCATC | AtSAND1 qPCR Primer |
| 12 | CACGTTTGGACCTTCAAGTATAGG | TCACCTAGCTGCAGACCATGA | AtFUS3 qPCR Primer |
| 13 | TGGACCAGCACAGCAACAAC | GTTGCTGCTGGACCACGATA | AtLEC1 qPCR Primer |
| 14 | CGGTATGACTGGATATGAAC | ATTTGCATCGGAGAGCTC | AtARR6 qPCR primer |
| 15 | CGAAGGGTTTAGGACTACATGAAG | GTGGGTTCACATGATGGTGCAA | AtCLV3 qPCR Primer |
